# NRH, a potent NAD+ booster, improves glucose homeostasis and lipid metabolism in diet-induced obese mice though an active adenosine kinase pathway

**DOI:** 10.1101/2024.08.29.610289

**Authors:** Xinliu Zeng, Yongjie Wang, Karina Farias, Andrew Rappa, Chirstine Darko, Anthony Sauve, Yue Yang

## Abstract

NAD^+^ deficiency underlies obesity-induced metabolic disturbances. Here we evaluated the treatment effect of a new and potent NAD^+^ enhancer, dihydronicotinamide riboside (NRH), in diet-induced obese mice with hyperglycemia and hyperlipidemia. Administering NRH for 7 weeks improved glucose homeostasis by enhancing pancreatic beta-cell functional mass, increasing muscle insulin sensitivity, and reducing hepatic gluconeogenesis. NRH treatment also mobilized fat deposition, reduced circulating lipid, and improved white adipose function. Significant elevation in multi-tissue NAD^+^ levels and sirtuin (SIRT) activities, especially SIRT3, mediated these metabolic improvements. Inhibiting adenosine kinase (ADK), a newly recognized enzyme in the NRH-induced NAD^+^ synthesis pathway, blocked NRH’s effect in improving glucose and lipid metabolism. ADK inhibition also reduced tissue NAD^+^ elevation and the subsequent activation of SIRT3, suggesting an active ADK pathway is necessary for NRH-induced metabolic benefits. These observations, for the first time, establish NRH as a promising intervention for correcting obesity-induced metabolic syndrome.

## Introduction

Rates of overweight and obesity have steadily increased across all population groups, leading to higher morbidity and mortality from cardiovascular diseases (CVD), type 2 diabetes (T2D), and other chronic illnesses, creating a significant burden in the healthcare systems^1^. However, managing obesity has been proven a challenging task. NAD^+^, a redox cofactor and a central metabolite for various physiological processes, offers a promising therapeutic opportunity to address obesity-related metabolic problems^2^. Decrease of blood and tissue NAD^+^ has been reported in elderly or obese humans and is associated with higher prevalence of metabolic diseases^3,4^. Previous studies have demonstrated that chronic high fat diet (HFD) feeding in mice causes NAD^+^ depletion in tissues such as the liver^5^, adipose^6^ and skeletal muscle^7^. NAD^+^ deficiency reduced the activity of sirtuins, which are NAD^+^-dependent Class III histone deacetylases and key regulators of energy metabolism, oxidative stress response, as well as glucose and lipid metabolisms^8^. Replenishing NAD^+^, either with inhibitors of major NAD^+^ consumers like cluster of differentiation 38 (CD38)^9^ or Poly (ADP-ribose) polymerase 1 (PARP1)^10^, or using precursors nicotinamide riboside (NR)^11–13^ and nicotinamide mononucleotide (NMN)^5,14,15^, has been shown to enhance sirtuin activities and protect mice against HFD-induced obesity and related metabolic syndrome. However, metabolite tracing revealed that the majority of NR or NMN was degraded to nicotinamide or nicotinic acid before delivered to target tissues for NAD^+^ synthesis, and their efficacy in boosting tissue NAD^+^ levels were limited^16–18^. This may account for the marginal improvements reported in many human trials with pre-diabetic or diabetic subjects^19,20^.

Our lab has previously synthesized a novel NAD^+^ precursor named dihydronicotinamide riboside (NRH)^21^. NRH is metabolized through a novel pathway, where adenosine kinase (ADK) phosphorylates NRH into NMNH, which is then either oxidized into NMN or converted to NADH for NAD^+^ synthesis^22^. NRH is more effective than NR and NMN in elevating NAD^+^ levels in cells and mouse tissues, increasing cellular NAD^+^ by up to 10-fold and boosting liver NAD^+^ by over 5-fold^21^, extents unmatched by any other NAD^+^ boosting treatments. However, the biological consequences of continuous high NAD^+^ induction from prolonged NRH treatment remain unclear. In this study, we investigated the safety and therapeutic effects against metabolic disorders of long-term NRH administration in mice that were lean or obese.

## Results

### NRH treatment has long-lasting effect in raising blood NAD^+^ and altered body composition in mice

Firstly, we examined the effectiveness and safety of NRH administration in CD-fed mice. Young, male C57BL/6J mice were intraperitoneally (IP) injected with vehicle or 250 mg/kg NRH, the lowest dose to maximally increase liver NAD^+^ levels^21^, 3 times a week for a continuous period of 7 weeks. Blood NAD^+^ levels were monitored to verify the treatment’s effectiveness. Two hours after NRH injection, blood NAD^+^ was increased by 1.5-fold, and stayed elevated until 48 hours (**S. Fig 1A**). Additionally, the blood NAD^+^ levels were measured at 72-hour post-injection, which showed over 1.5-fold increase, twice during the treatment period (**S. Fig 1B**). No differences in food consumption (data not shown) or body weight (**S. Fig 1C**) were observed throughout the 7 weeks between the CD control and NRH groups. Echo-MRI analysis in the 6th week revealed a slight increase in lean mass and a reduction in fat mass in the NRH group (**S. Fig 1D**). After sacrifice, serum ALT and AST levels were measured, showing no differences between groups (**S. Fig 1E**), suggesting that this dose and treatment regimen were well tolerated by the mice and did not cause overt toxicity.

We then investigated whether NRH has any therapeutic effect in mice with established obesity and related metabolic disorders. Similar NRH treatment regimen was administered to DIO mice that had been fed with HFD for 3 months (**Fig 1A**). Blood NAD^+^ levels were examined over the first 7 days, showing the first two doses significantly increased in blood NAD^+^, whereas the third dose has reduced effect, but the baseline levels were elevated throughout the period (**Fig 1B**). Afterwards, the baseline blood NAD^+^ levels were maintained at an average increase of 1.8 to 1.9-fold above control, demonstrating NRH’s overall effectiveness throughout the study (**Fig 1C**).

During the 7-week period, no differences in food consumption (**Fig 1D**) or average body weight (**Fig 1E**) were observed between the HFD control and NRH groups. However, the total body weight gain was significantly lower in the NRH-treated group (**Fig 1F**). Serum ALT and AST measured after sacrifice also revealed no differences between groups (**Fig 1G**). Furthermore, Echo-MRI showed similar changes in body composition as in the CD groups, with a slight but significant increase in lean mass and a trend towards reduced fat mass in the NRH-treated mice (**Fig 1H**). To determine if the changes in body weight gain and body composition were due to elevated energy expenditure, we monitored the respiration and baseline activities in these mice using metabolic cages. No differences in CO_2_ production, oxygen consumption, whole-body respiratory exchange ratio (RER) or food intake (**Fig 1I**) were observed between groups. There was also no significant difference in the distance traveled (**Fig 1J**), indicating similar levels of physical activity in mice with or without NRH treatment. Therefore, the changes in NRH-induced body composition were not caused by elevated energy expenditure.

**Figure 1.**
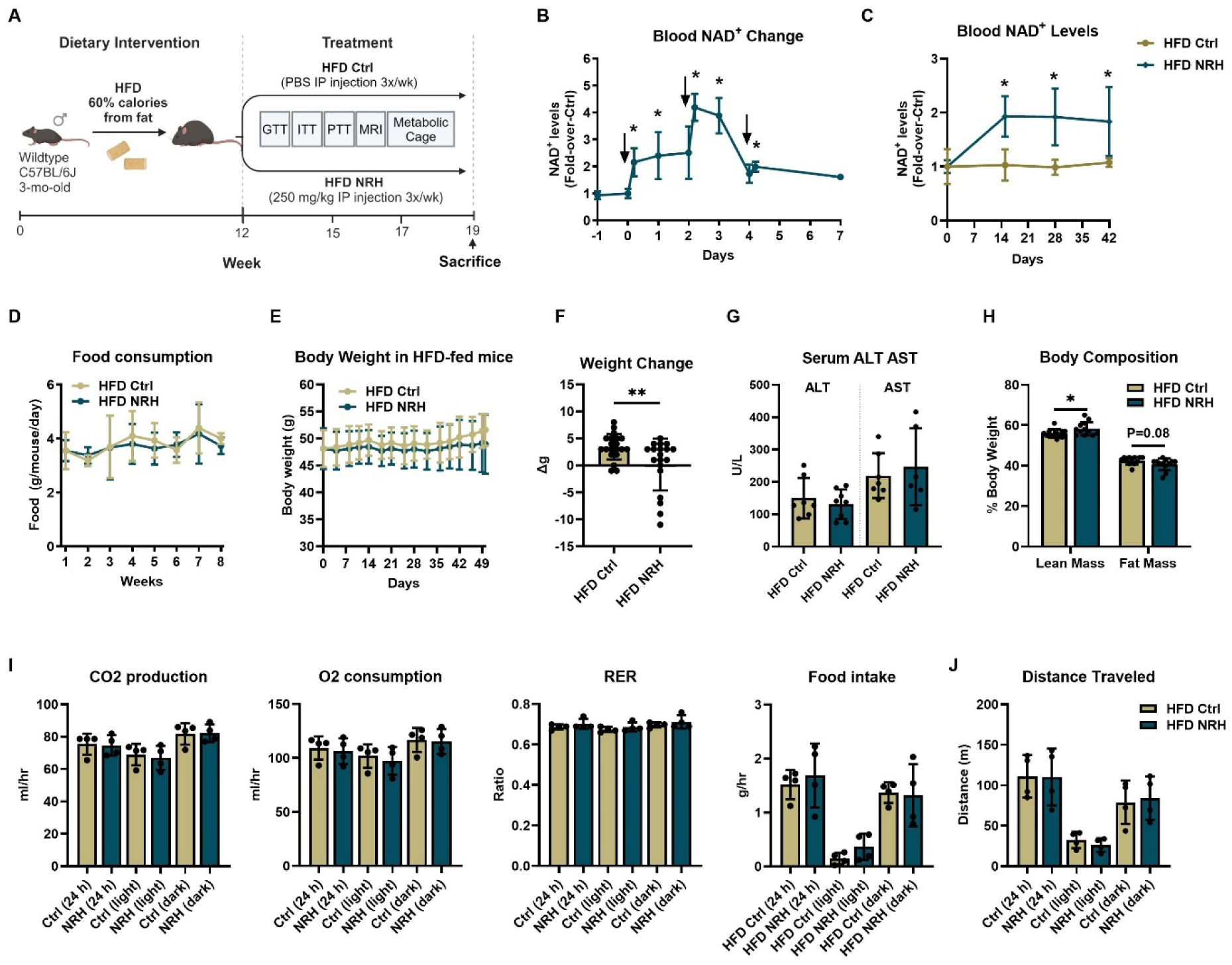
NRH treatment has a long-lasting effect in raising blood NAD^+^ and altering body composition in mice. A) Experiment scheme in HFD-fed mice with 20 mice per group. B) Changes in blood NAD^+^ levels over the first 7 days. Arrows indicate NRH injection. C) Baseline blood NAD^+^ levels over the 7-week treatment period. D) Food consumption over the treatment period. E) Changes in average body weights over the 7 weeks. N=20 per group. F) Total body weight change before and after the treatment. N=20 per group. G) Serum ALT and AST levels of HFD-fed mice at sacrifice. H) Body composition measurement in HFD-fed mice at the 5th week of treatment. N=7-8 per group. I) HFD-fed mice were monitored with a metabolic cage at the 5th week of treatment. Their oxygen consumption, CO_2_ production, respiratory exchange ratio (RER), and food intake with 12 hours of light and 12 hours of dark were recorded. J) Distance traveled was recorded over 24 hours with 12 hours of light and 12 hours of dark phases. Unless specified, N=5 per group. Data are shown as mean ± SD. * indicates p<0.05 and ** indicates p<0.01 compared to time 0 or HFD Ctrl.

### NRH improved glucose homeostasis in HFD-fed mice

HFD feeding in mice is known to induce hyperglycemia and insulin resistance^23^. Preliminary glucose tolerance test (GTT) and insulin tolerance test (ITT) demonstrated that the DIO mice had already developed glucose intolerance and insulin resistance before any treatment (**S. Fig 2A, B**). After 3 weeks of NRH administration, we assessed glucose homeostasis in these mice. NRH-treated mice exhibited significantly faster glucose clearance rates and lower area-under-the-curve (AUC) following a bolus dextrose injection compared to the controls (**Fig 2A**). Fasting glucose levels in the NRH group were reduced by almost half (**Fig 2B**) and were similar to those in the lean control group (**S. Fig 2C**). To determine whether this improvement was due to enhanced insulin secretion, glucose-stimulated insulin secretion (GSIS) was evaluated. NRH-treated group secreted more insulin into the serum in the first 30 minutes following dextrose injection (**Fig 2C**), despite having similar fasting insulin levels (**Fig 2D**), demonstrating that NRH treatment improved the functionality of pancreatic beta cells. To further investigate if the functional beta cell mass had improved, we quantified total insulin content using acidic-ethanol extracts of the pancreas after sacrifice and found a significant increase in the NRH-treated group (**Fig 2E**). Immunohistochemistry (IHC) staining confirmed this, showing more insulin present in the pancreas slides (**Fig 2F**) and the ratiometric quantification revealed an increase in insulin-stained area and beta cell mass (**S. Fig 2D**). These results suggest a pancreas-specific corrective effect induced by NRH treatment, which rescued both function and mass of beta cells to combat obesity-induced beta cell damage. In contrast, in lean mice, their glucose clearance rates were unchanged by NRH treatment (**S. Fig 2E**). There was also no difference in their fasting glucose and insulin levels (**S. Fig 2E-F**). However, an increase in total pancreatic insulin content in the NRH treated lean mice was observed, echoing the increase in beta cell mass seen in DIO mice (**S. Fig 2G**).

**Figure 2.**
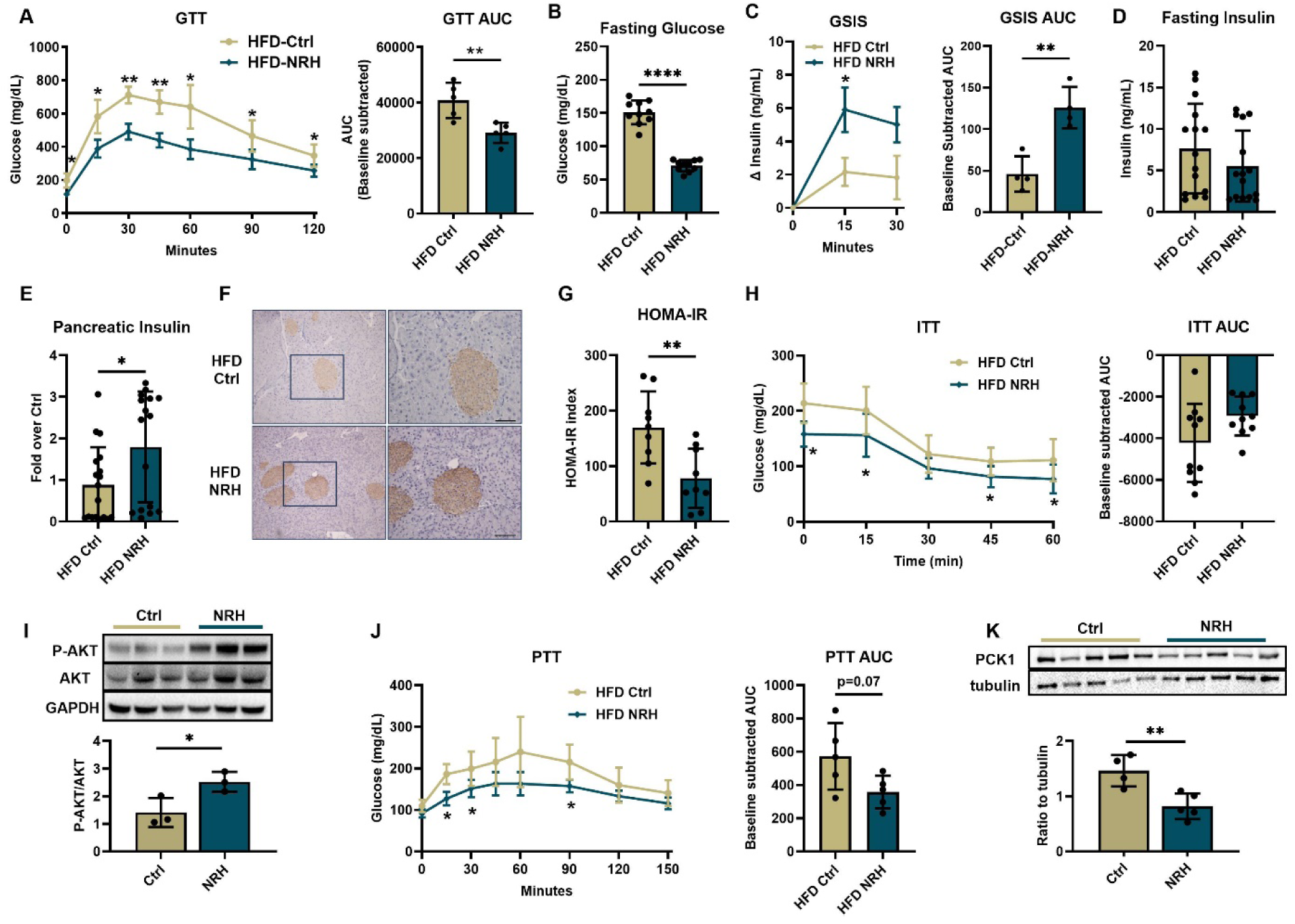
NRH significantly improved glucose homeostasis in HFD-fed mice. A) GTT in HFD-fed mice at the 3rd week of treatment. Blood glucose was measured after a bolus of glucose injection at multiple time points following a 16-hour fasting. Baseline-subtracted area-under-the-curve (AUC) was calculated based on GTT. N=10 per group. B) Blood glucose levels after fasting. C) Serum insulin changes and AUC during the first 30 minutes of GTT. N= 4 per group. D) Serum insulin levels after 16-hour fasting. E) Total pancreatic insulin levels measured after sacrifice. N= 20 per group. F) Representative IHC of pancreas staining for insulin. Images were taken under 10x and 20x magnification. G) HOMA-IR calculated based on fasting glucose and fasting insulin levels. H) ITT in HFD-fed mice at the 6th week of treatment after 4-hour fasting. Blood glucose was measured after insulin IP injection at 15-minute intervals. Baseline-subtracted AUC was calculated. N =10 per group. I) Protein expression of p-AKT, AKT, and GAPDH in HFD-fed mice muscle. p-AKT and AKT protein levels were quantified and presented as ratios. J) PTT and its AUC in HFD-fed mice at the 6th week of treatment. Blood glucose was measured after pyruvate injection at multiple time points. N=5 per group. K) Protein expression of PCK1 and tubulin, and their ratio in mice liver. Data are shown as mean ± SD. * indicates p<0.05, ** indicates p<0.01, **** indicates p<0.0001 compared to HFD Ctrl.

Because the Homeostasis Model Assessment of Insulin Resistance (HOMA-IR) predicted a reduction in insulin resistance within NRH-treated DIO mice (**Fig 2G)**, we performed an ITT to test whole-body insulin sensitivity. The NRH-treated group exhibited lowered glucose levels at multiple time points, suggesting an improvement in insulin sensitivity, despite no changes in baseline subtracted AUC were observed (**Fig 2H**). Post-mortem analysis revealed elevated p-AKT/AKT levels, an indicator of activated insulin signaling, within the skeletal muscle of NRH-treated mice (**Fig 2I**), further confirming their insulin sensitivity has been improved. The enhanced insulin sensitivity induced by NRH was observed only in mice with pre-existing insulin resistance, and had no effect in healthy, lean mice (**S. Fig 2H**).

Endogenous glucose production, another contributor to hyperglycemia in mice, is potentially suppressed by NRH treatment, leading to reduced fasting glucose in blood (**Fig 2B**). To assess the gluconeogenesis rate, we IP injected HFD-fed mice with pyruvate as a substrate to induce glucose production. The NRH group displayed lowered blood glucose at multiple time points, and smaller AUCs than the control group (**Fig 2J**). To validate the suppression of the gluconeogenic pathway, we measured the hepatic protein level of phosphoenolpyruvate carboxykinase 1 (PCK1), a rate-limiting enzyme in gluconeogenesis, and found it significantly reduced in NRH group (**Fig 2K**). Together, these results demonstrate that NRH works through multiple tissues and mechanisms to improve glucose homeostasis in obese mice.

### NRH ameliorated hyperlipidemia by mobilizing fat deposition in DIO mice

Hypercholesterolemia and hypertriglyceridemia are obesity-induced risk factors contributing to the pathogenesis and progression of cardiovascular diseases^24^. Niacin, or nicotinic acid, a metabolite increased following NRH administration^22^, is commonly used as a treatment for dyslipidemia^25^. To examine if NRH treatment can alter lipid metabolism in mice, we measured fasting cholesterol and triglyceride (TG) levels in serum. In HFD-fed mice, total cholesterol (TC), as well as high-density lipoproteins cholesterol (HDL-C) and low-density lipoproteins cholesterol (LDL-C), were all significantly reduced by NRH (**Fig 3A**). Fasting TG (**Fig 3B**), but not free fatty acid (FFA) (**Fig 3C**), was also lowered. Blood analysis also revealed that NRH reduced oxidative stress in the blood, as indicated by lower levels of malondialdehyde (MDA), a product of lipid peroxidation (**Fig 3D**). There was no significant change in inflammatory status represented by interleukin 6 (IL6) levels (**Fig 3E**). In lean mice, TC, TG, FFA and MDA levels did not differ between groups (**S. Fig 3A-D**), showing the hypolipidemic effect of NRH occur only in obese mice.

**Figure 3.**
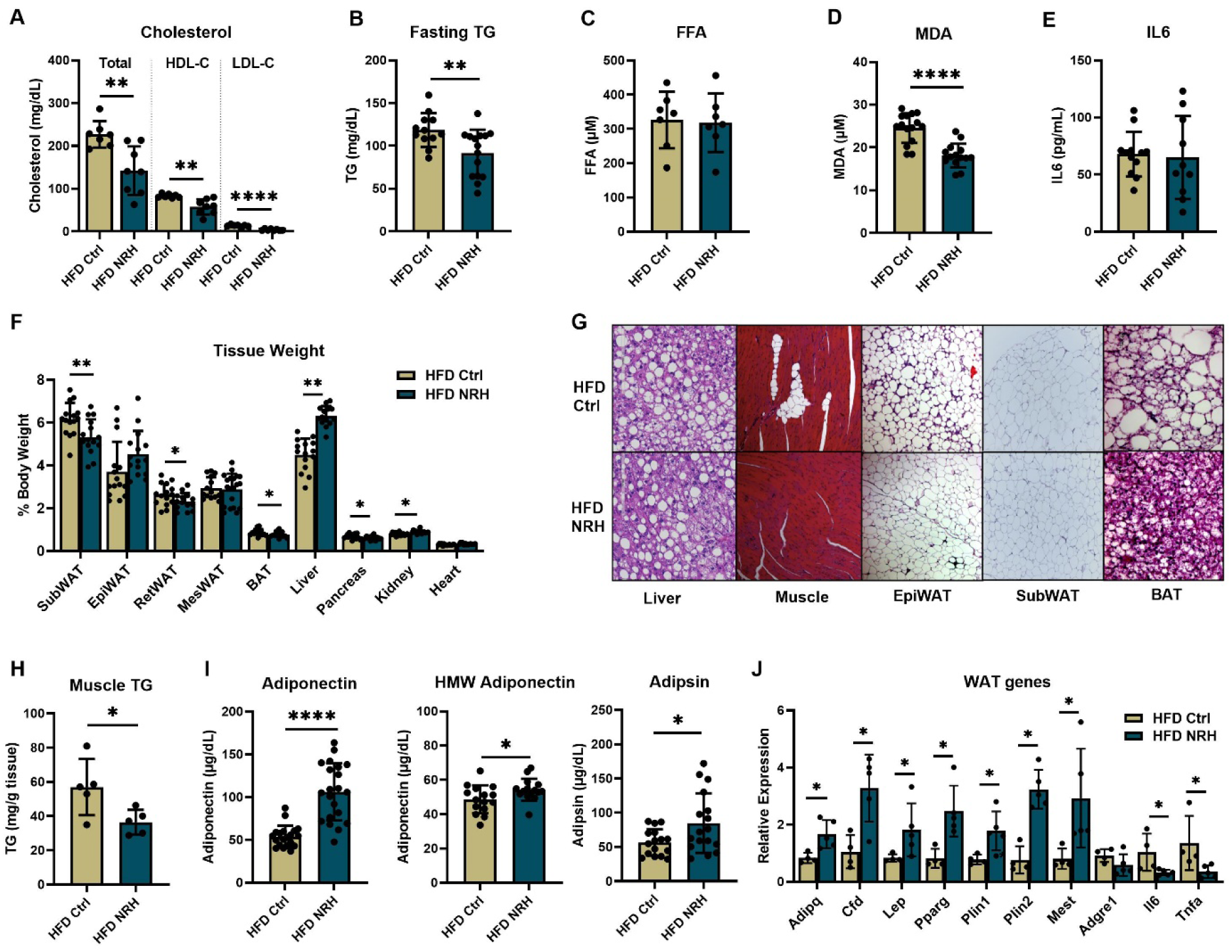
NRH ameliorated hyperlipidemia by mobilizing fat storage in DIO mice. A) Serum TC, HDL-C and LDL-C in 16-hour fasted, HFD-fed mice. N=7-8. B) Serum TG levels in fasted mice. N=10-15. C) Serum FFA in mice. N=7. D) Serum MDA levels in mice. N=15. E) Serum IL6 levels in mice. N=10-12. F) Tissue weights shown in percentage of body weight between HFD groups. N=15. G) Representative H&E staining image of liver, muscle, epidydimal WAT, subcutaneous WAT and BAT in HFD mice with or without NRH treatment. H) Tissue TG content in muscle. N=5. I) Serum adipokines, including total adiponectin, HMW adiponectin and adipsin. N=15. J) Gene expressions on adipokines, lipid metabolism, cell proliferation, and inflammation markers in subcutaneous WAT normalized to 18S expression. N=5. Data are shown as mean ± SD. * indicates p<0.05, ** indicates p<0.01, **** indicates p<0.0001 compared to HFD Ctrl.

To further corroborate if NRH altered body composition as indicated by Echo-MRI, tissues were weighed after sacrifice. In lean mice, there were no significant changes in tissue weights even though trends were observed (**S. Fig 3E**). Tissue TG contents were similar between groups (**S. Fig 3F**). However, in DIO mice, weights of subcutaneous and retroperitoneal white adipose tissue (WAT) and brown adipose tissue (BAT) were significantly reduced in the NRH group (**Fig 3F**). H&E staining showed the BAT from the NRH group exhibited smaller fat droplets, implying the reduced BAT weight resulted from less fat deposition (**Fig 3G**). Epididymal WAT, with unchanged weight, exhibited fewer signs of inflammation represented by crown-like structures, whereas subcutaneous WAT, showed no apparent differences in morphology (**Fig 3G**). Additionally, a significant amount of ectopic fat deposition was found in the skeletal muscle of HFD control mice but was reduced by NRH treatment (**Fig 3G**). In support of this finding, lower average TG concentration in the muscle of the NRH group were observed (**Fig 3H**).

To examine if the changes in WAT weight are associated with improvement in their functions, circulating levels of adipokines were measured. The presence of insulin sensitizing adipokines adipsin^26^ and adiponectin, especially high-molecular-weight (HMW) adiponectin^27^, were all significantly higher in NRH-treated DIO mice (**Fig 3I**), but were unchanged in lean mice (**S. Fig 3G**). Follow-up analyses demonstrated corresponding elevations in mRNA expression of *Adipoq* (adiponectin), *Cfd* (adipsin)*, Lep, Pparg, Plin1, Plin2,* and *Mest* in the subcutaneous WAT, accompanied by reduced inflammatory markers *Il6* and *Tnfa* (**Fig 3J**). These data demonstrated that NRH treatment reduced lipid deposition in WAT and BAT while improving adipose function by facilitating adipokines secretion in obese mice.

Conversely, we found NRH treatment increased the liver weights of DIO mice by an average of 20% (**Fig 3F**), while no difference in liver weights were seen in lean mice (**S. Fig 3E**). H&E staining revealed similar levels of steatosis in the livers of both HFD control and NRH groups with no overt liver damage (**Fig 3G**), which was supported by similar ALT and AST levels in the blood (**Fig 1G**). We then compared the liver TG contents and found similar TG concentrations (**Fig 4A**). Lipidomic analysis revealed a shift from shorter to longer TG species in NRH-treated group (**Fig 4B**).

**Figure 4.**
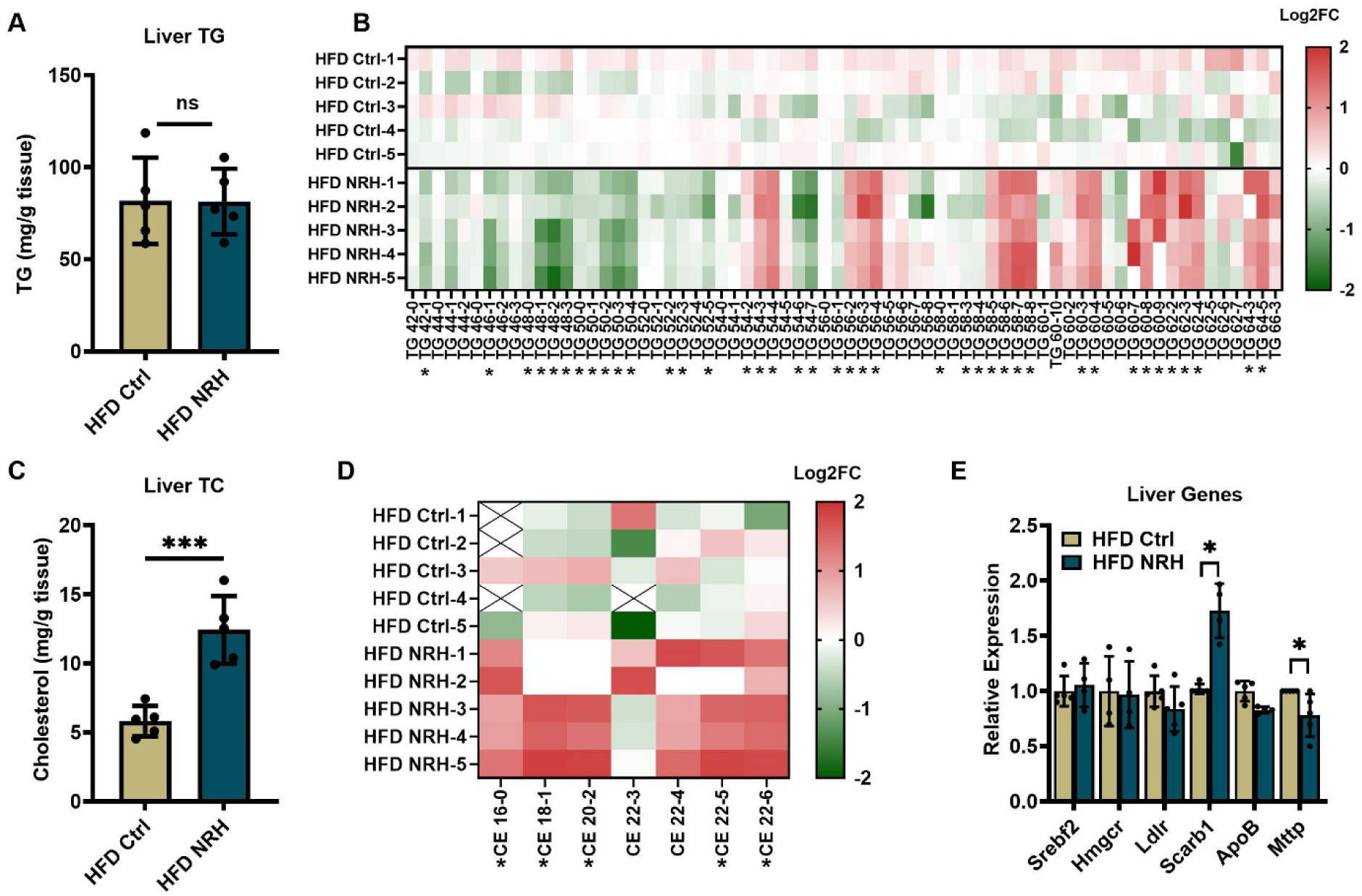
NRH treatment altered lipid species in the liver of DIO mice. A) Liver TG levels in HFD-fed mice. B) Heatmap of different TG species extracted from HFD Ctrl and HFD NRH livers. The color represents Log2FC over average control values. * marks adjusted p<0.05. C) Liver TC levels in HFD-fed mice. D) Heatmap of CE species in HFD-fed mice. The color represents Log2FC over average control values. * marks adjusted p<0.05. E) mRNA analysis showing differential expression in genes related with cholesterol metabolism secretion normalized to 18S expression from liver in NRH-treated mice. N=5 per group. Data are shown as mean±SD, * indicates p<0.05 compared to HFD Ctrl.

Cholesterol is another major component of lipid droplets. We then quantified the TC levels in the liver, and found NRH-treated livers had 2-3 times more TC than those of HFD controls (**Fig. 4C**). Compared to free cholesterol, cholesteryl ester (CE) is considered a neutral, benign form for storage^28^. Lipidomics showed NRH increased in almost all CE species (**Fig 4D**). We infer that the increased CE storage may be responsible for the lower cholesterol levels in circulation. To examine the cholesterol metabolism pathway, we analyzed the mRNA expression of key genes. Although no significant differences were found in cholesterol synthesis or cholesterol ester synthesis genes such as *Srebp2*, *Hmgr* and *Ldlr*, we observed an elevation in the expression of Scavenger Receptor Class B Type 1 (*Scarb1*), a receptor involved in HDL-C uptake from circulation^29^, and a suppression in the expression of microsomal triglyceride transfer protein (*Mttp*), a gene responsible for the production of VLDL-C released into circulation from the liver^30^ (**Fig 4E**). Thus, NRH treatment may increase the uptake of HDL-C while reducing the release of VLDL-C, resulting in the accumulation of CE in the liver and reduced TC levels in circulation. This could potentially reduce risks of metabolic disorders associated with hyperlipidemia despite the observed increase in liver weights. Overall, the observations confirmed a mobilization of lipid storage in the body, which led to improved adipose function, less ectopic TG deposition, with the exception of an increase in liver TG and CE storage.

### NRH increased in NAD^+^ levels and activated sirtuins in tissues

We found the HFD-feeding significantly reduced NAD^+^ levels in the blood, liver, pancreas, heart, subcutaneous WAT and BAT comparing to CD-feeding (**Fig 5A**), consistent with previous publications showing that a depleted NAD^+^ pool underlies obesity-induced disorders. We hypothesize that as an NAD^+^ precursor, NRH achieves its beneficial effects through elevating tissue NAD^+^ levels. Following the last treatment, tissues were collected after either 4 or 24 hours to assess their NAD^+^ content. In lean mice, only the pancreas showed increased NAD^+^ levels at 24 hours post NRH injection (**S. Fig 4**). In obese mice, NAD^+^ levels in the liver, pancreas, kidney, heart, and subcutaneous WAT peaked at 4 hours and then returned to baseline by 24 hours. Blood, muscle, and BAT maintained an elevated NAD^+^ level from 4 to 24 hours (**Fig 5B**). These findings indicate that the ability of NRH to induce NAD^+^ elevation is ubiquitous and potent; however, the NAD^+^ increases induced by NRH are relatively transient in tissues compared to blood.

**Figure 5.**
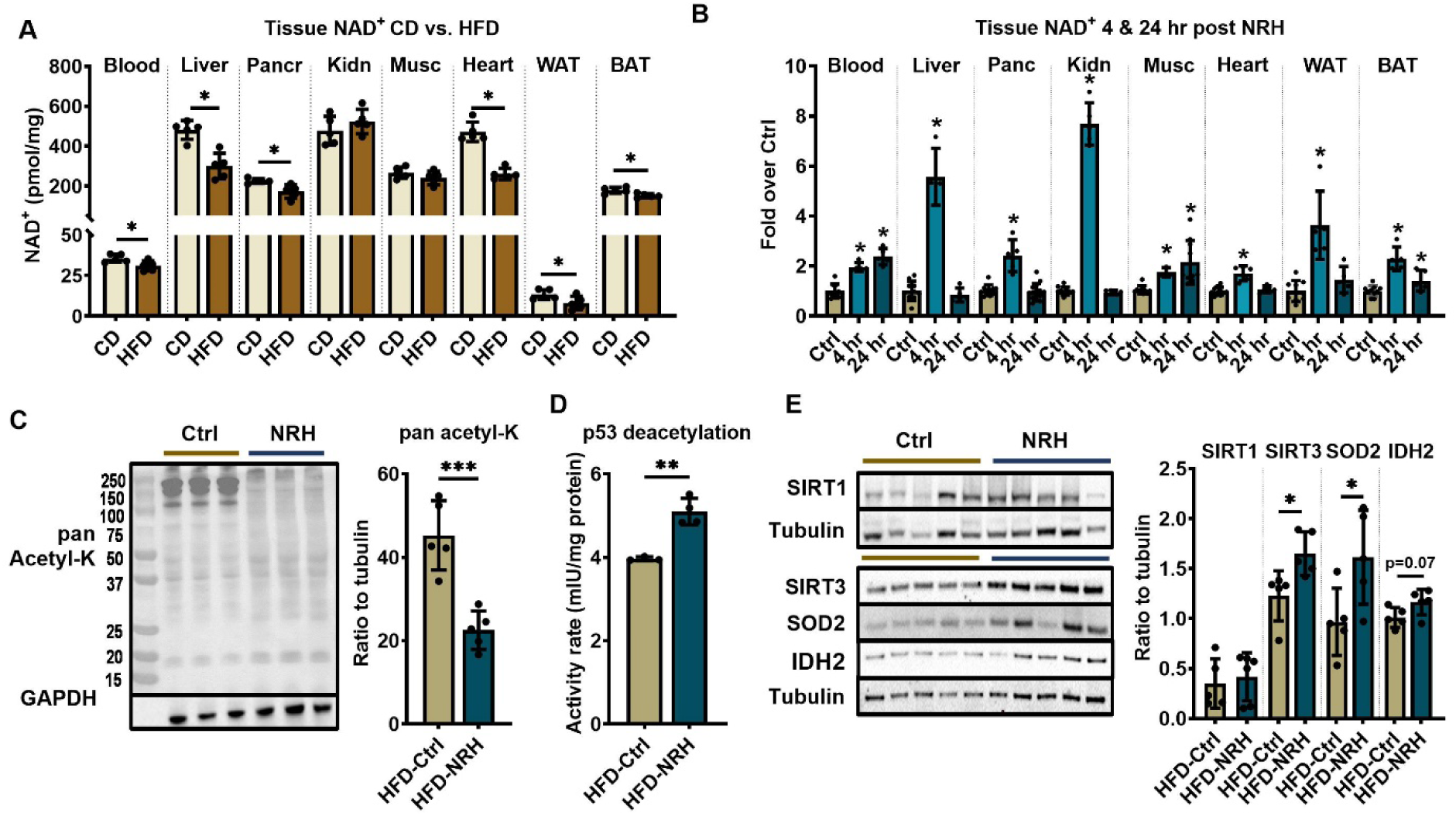
NRH boosted up tissue NAD^+^ levels and activated sirtuins. A) Tissue NAD^+^ levels between CD- and HFD-fed Ctrl mice. B) NAD^+^ levels in tissues 4- and 24-hour after the final NRH injection in the HFD-fed mice compared to HFD Ctrl. N= 5-15 per group. C) Pan acetyl-lysine in liver protein lysates from HFD-fed mice 4-hour post injection and its quantification over GAPDH expression. D) Deacetylation activities of liver lysates 4-hour post injection with acetyl-p53 as substrate. N=4. E) Protein expression and quantification of SIRT1, SIRT3 and downstream target SOD2 and IDH2 in liver lysates with tubulin as loading controls. Each lane represents a single mouse. Data are shown as mean ± SD, * indicates p<0.05, ** indicates p<0.01, **** indicates p<0.0001 compared to HFD Ctrl.

To examine if the increase in NAD^+^ levels lead to activation of sirtuins, which are NAD^+^-dependent deacetylases, we analyzed the pan-acetyl lysine levels in liver protein lysates. The NRH-treated group showed significantly lower acetyl-lysine bands compared to the controls, indicating increased overall deacetylase activity (**Fig 5C**). This was supported by enhanced deacetylation of acetyl-p53, a common substrate for SIRT1, 2 and 3, following NRH treatment (**Fig 5D**). Next, we examined the protein expression of major sirtuin isoforms. NRH treatment led to higher expression of SIRT3, a mitochondrial isoform, in NRH-treated liver; however, the expression of SIRT1, a nuclear and cytosol isoform, was unchanged (**Fig 5E**). Mitochondrial NAD^+^ pool is the most stable among subcellular compartments and crucial for maintaining cellular metabolism and energy production^31^. Increased SIRT3 protein levels may indicate that the NRH-induced NAD^+^ enhancement sustained longer in mitochondria than in nucleus and cytosol, as evidenced by unchanged SIRT1 expression. Additionally, two mitochondrial proteins, superoxide dismutase 2 (SOD2) and mitochondrial isocitrate dehydrogenase (IDH2), are both targets for SIRT3-induced deacetylation. They are also enhanced transcriptionally through activation of Forkhead Box O (FOXOs) by SIRT1^32^ or SIRT3^33^-induced deacetylation. The protein expressions of both SOD2 and IDH2 were increased in the livers of NRH-treated mice, supporting the elevated sirtuin activities and improved mitochondrial antioxidant defense to combat excessive oxidative stress (**Fig 5E**). Overall, these data suggest that the benefits of NRH were mediated, at least partially, through the activation of sirtuins, particularly through SIRT3.

### Inhibition of ADK blocked the NRH-induced improvement in glucose and lipid metabolism

Previously, we identified ADK as the key enzyme in the NRH downstream pathway of NAD^+^ synthesis, phosphorylating NRH into NMNH for NAD^+^ synthesis^22^. However, there was also a marked increase of NR levels in the mouse liver, suggesting that some NRH may also be oxidized into NR *in vivo*, which later joins the NAD^+^ synthesis pathway.^22^. Therefore, whether ADK is indispensable in mediating the metabolic benefits induced by NRH remains unknown. ADK is an ubiquitously expressed enzyme that plays an essential role in producing AMP^34^. ADK deficiency leads to severe consequences, such as neonatal hepatic steatosis^35^, cerebrovascular abnormalities^36^, and other neurological and hepatic impairments^37^. To assess the necessity of intact ADK function in mediating NRH effects and to avoid undesirable side effects caused by prolonged ADK deficiency, we used a temporary pharmacological inhibitor of ADK, ABT702, which was administered 1 hour prior to each NRH injection in obese mice over a 7-week treatment period (**Fig 6A**). While no differences in average weight were observed between the groups (**S. Fig 5A**), weight gains were decreased by NRH treatment, with or without ABT702 (**Fig 6B**). Blood NAD^+^ levels were measured throughout the treatment period, and the NRH-induced elevations in NAD^+^ baseline were blocked by co-treatment with ABT702, demonstrating its effective inhibition of NRH metabolism (**Fig 6C**).

**Figure 6.**
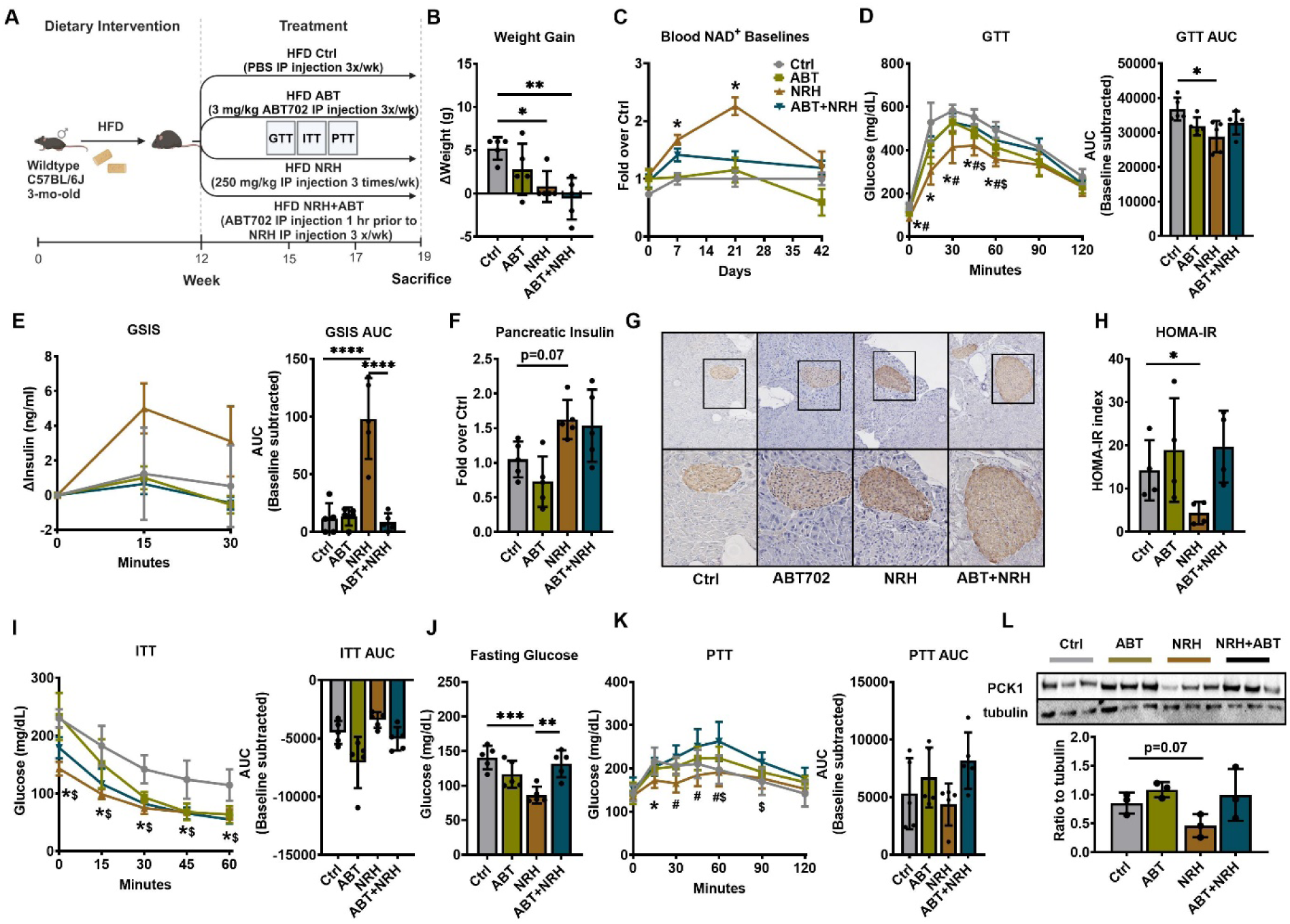
Inhibition of ADK blocked the NRH-induced improvement in glucose metabolism. A) Experiment scheme in HFD-fed mice treated with NRH, ABT702, or the combination of both, with 5 mice per group. B) Total body weight changes over the treatment period. C) Baseline blood NAD^+^ level throughout the 7-week treatment period in each group. D) Blood glucose levels during GTT and AUCs of GTT. E) Serum insulin level changes during the first 30 min of GTT and their AUCs. F) Total pancreatic insulin quantification through acidic-ethanol extraction. G) IHC of insulin in pancreatic islets. H) HOMA-IR calculated based on fasting glucose and fasting insulin levels. I) Blood glucose levels during ITT after 4-hour fasting and their AUCs. J) Blood glucose levels after 16-hour fasting. K) Blood glucose levels during PTT and the AUCs for PTT. L) Protein expressions of PCK1 in liver and their quantification. N=3-5 per group. Data are shown as mean ± SD, * indicates p<0.05 between Ctrl and NRH unless specified, ** indicates p<0.01, *** indicates p<0.001, **** indicates p<0.0001. ^#^ indicates p<0.05 between NRH and ABT+NRH, ^$^ indicates p<0.05 between Ctrl and ABT+NRH.

GTT analysis revealed that the significantly improved glucose clearance rates by NRH were abolished by ADK inhibition (**Fig 6D**). ABT702 blocked the improvement in insulin secretion induced by NRH (**Fig 6E**). However, increased pancreatic insulin contents were found in both NRH and NRH+ABT702 groups, with either acidic-ethanol extract (**Fig 6F**) or insulin staining (**Fig 6G**). Quantification of beta cell mass also showed a higher percentage of beta cells in the NRH+ABT702 group than NRH alone (**S. Fig 5B**), which may be due to the beta cell replication enhancing effect of ABT702 reported previously^38^, yet this increase in beta cell mass did not improve its function in insulin secretion.

Although HOMA-IR predicted more insulin resistance (**Fig 6H**), the NRH+ABT702 group exhibited comparable improvement in insulin sensitivity, showing similar performance as the NRH group during ITT (**Fig 6I**). Moreover, NRH-induced reductions in fasting glucose were reversed by ADK inhibition (**Fig 6J**). PTT suggested that the suppressed gluconeogenesis by NRH treatment was negated by ADK inhibition (**Fig 6K**), and the reduced protein expression of PCK1 was also restored by co-administration of ABT702 (**Fig 6L**). Together, the use of an ADK inhibitor blocked the majority of NRH-induced improvements in glucose homeostasis, with the exception of enhanced beta cell mass and insulin sensitivity, indicating that intact ADK function is crucial for mediating the metabolic benefits of NRH treatment.

To test if NRH-induced improvements in lipid metabolism are also mediated through ADK, we measured fasting TC and TG levels in serum. Co-administration of NRH with ABT702 reversed the reduction in TC observed with NRH alone (**Fig 7A**). However, fasting TG remained reduced in both the NRH and NRH+ABT702 groups, with the ABT702-only group also showing a trend towards lower fasting TG, suggesting a potential synergistic effect of NRH and ABT702 treatment on TG levels (**Fig 7B**). The NRH-induced improvements in adiponectin and adipsin secretion were blocked by co-treatment with the ADK inhibitor (**Fig 7C**). Further gene expression analysis revealed that NRH-induced increases in *Adipq, Cfd, Lep, Pparg, Plin1,* and *Plin2* in subcutaneous WAT were reversed, while the suppressions of inflammatory markers *Il6* and *Tnfa* by NRH were overturned by ABT702 co-treatment (**Fig 7D**).

**Figure 7.**
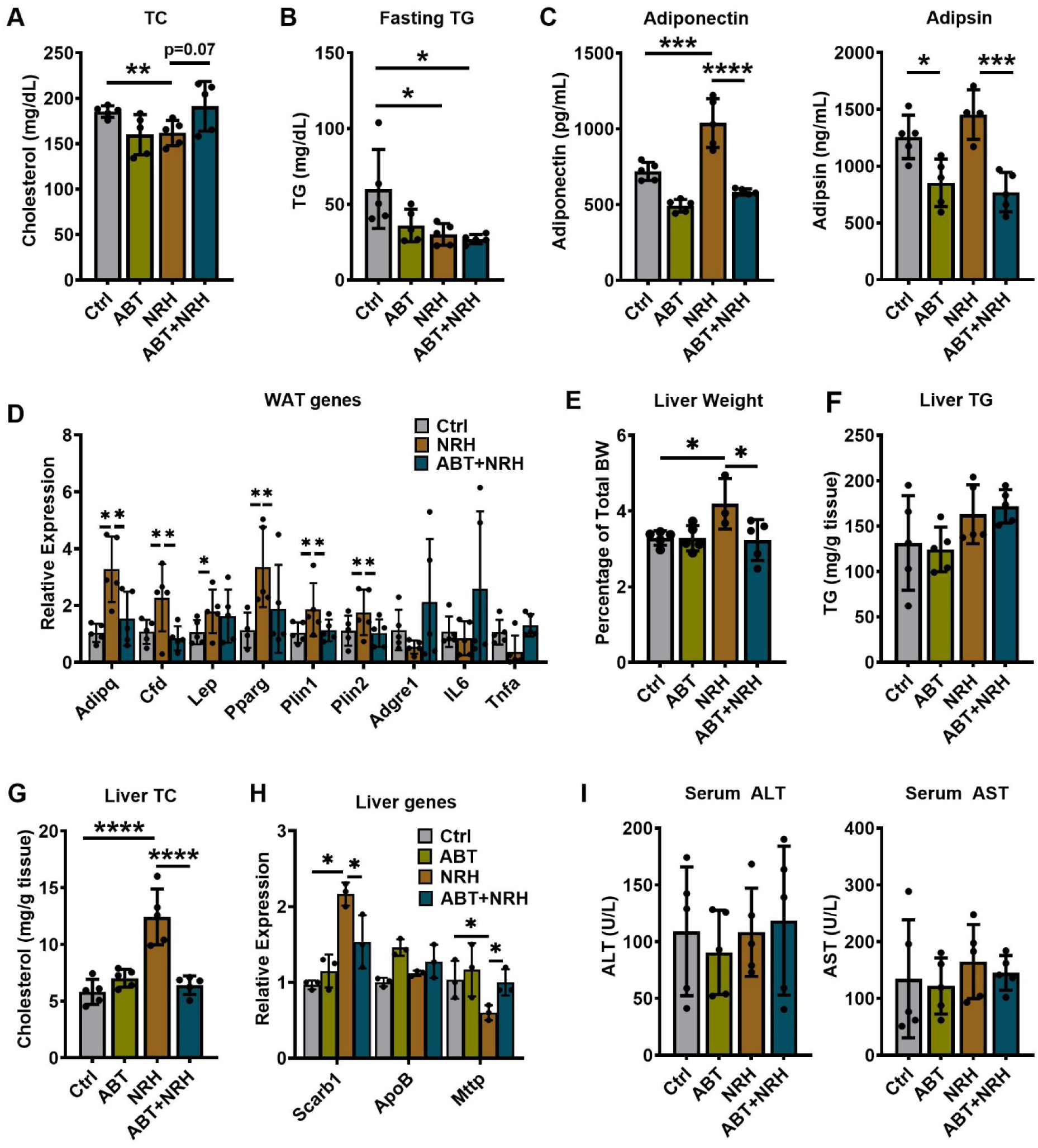
ADK inhibition prevented NRH-induced WAT improvement and lipid metabolism changes. A) Serum TC levels after 16-hour fasting. B) Serum TG levels after 16-hour fasting. C) Serum adiponectin and adipsin levels. D) Gene expressions on adipokines, lipid metabolism, cell proliferation, and inflammation markers normalized to 18S expression in subcutaneous WAT. E) Liver weights shown in percentage of body weight in HFD-fed mice with different treatment. F) Liver TG concentrations. G) Liver CE concentrations. H) mRNA analysis showing differential expression in genes related with cholesterol metabolism normalized to 18S expression in liver. I) Serum ALT and AST levels of HFD-fed mice at sacrifice. Data are shown as mean±SD, * indicates p<0.05, ** indicates p<0.01, *** indicates p<0.001, **** indicates p<0.0001.

NRH treatment increased liver weight, but this effect was removed in the NRH+ABT702 group, in which liver weight returned to control levels (**Fig 7E**). ADK inhibition blocked the NRH-induced increase in hepatic TC content (**Fig 7G**) but did not change their TG concentrations (**Fig 7F**). Gene analysis of cholesterol metabolism-related genes showed the NRH-induced changes in *Scarb1* and *Mttp* expressions reversed with ABT702 treatment (**Fig 7H**). Again, no differences were found in the serum ALT and AST levels between groups (**Fig 7I**), showing no apparent liver and tissue damage. Therefore, without the full activity of ADK, NRH did not stimulate CE accumulation via enhanced uptake and reduced release of cholesterol from circulation. Consequently, NRH did not reduce circulating cholesterol levels and did not increase liver weight in the presence of ADK inhibitor.

### AK inhibition partially inhibit NRH-induced NAD^+^ elevation and Sirt3 enhancement

Due to significant changes in NAD^+^ levels observed in multiple tissues 4 hours post-NRH injection, we analyzed tissue NAD^+^ changes 4 hours after the final injection within the 4 groups. NRH-induced NAD^+^ induction was significantly reduced, if not fully blocked, by co-administration of ABT702 in the blood, liver, pancreas, kidney, muscle, and BAT, indicating effective inhibition of NRH-induced synthesis (**Fig 8A**). The remaining increase in NAD^+^ levels is likely attributed to the conversion of NRH into NR or NAM. We also assessed whether sirtuin activities were altered. Pan-acetyl lysine analysis of liver protein lysates showed reduced acetylated proteins in both NRH and NRH+ABT702 groups, with no significant differences observed with or without ABT702 administration (**Fig 8B**). However, the increased deacetylation activity on acetyl-p53 in NRH group was reduced when combined with ABT702 co-treatment (**Fig 8C**). The NRH-enhanced SIRT3 protein expression was blocked by ABT702 co-treatment (**Fig 8D**), while ABT702 or in combination with NRH did not alter SIRT1 expression (**S. Fig 5C**). Additionally, downstream target expression of SOD2 and IDH2 proteins was reduced with ADK inhibition (**Fig 8D**), implying that sirtuin-mediated downstream effects contribute to the NRH-induced improvements, which were ameliorated by ADK blockage.

**Figure 8.**
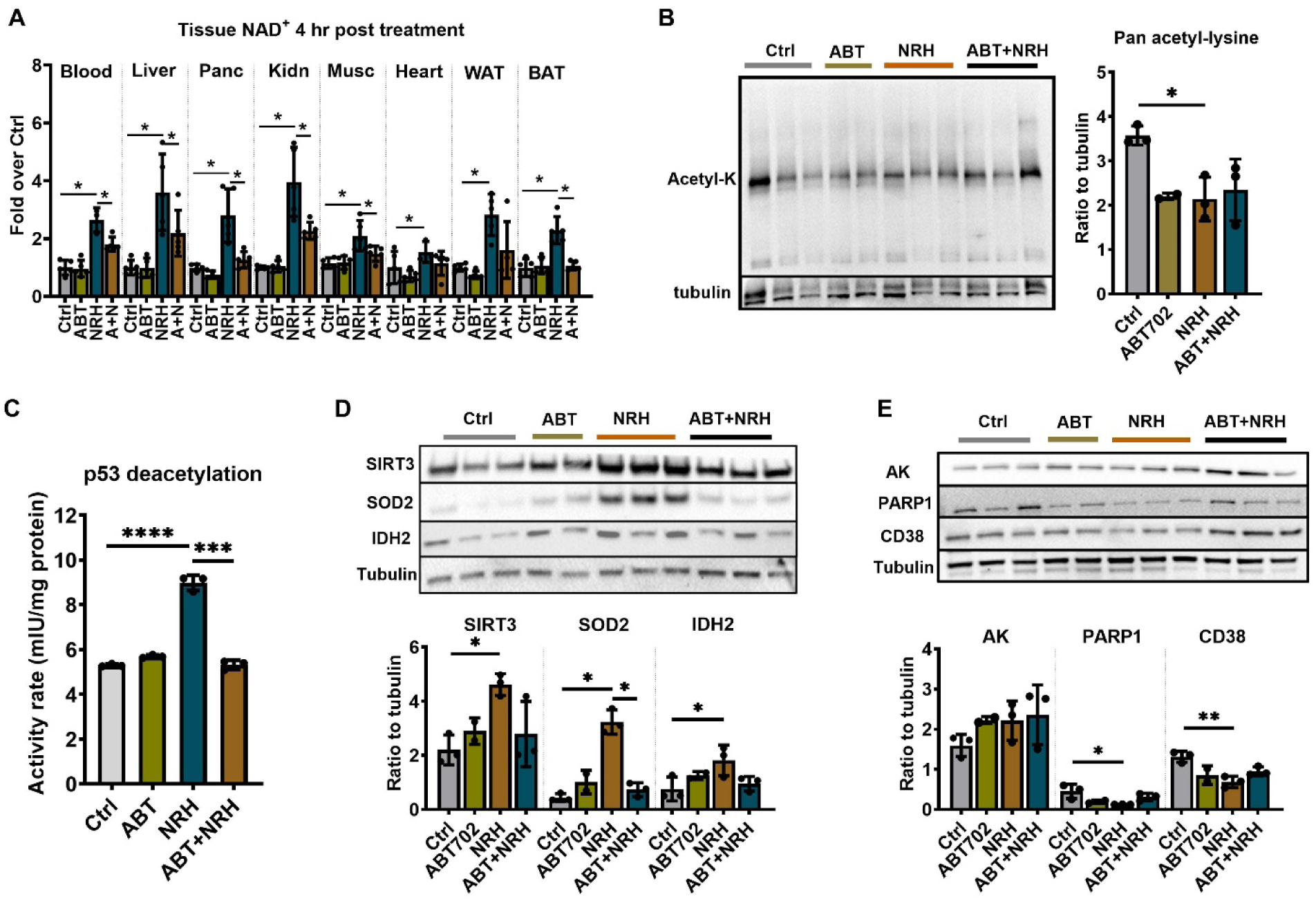
ADK inhibition partially blocked NRH-induced NAD^+^ elevation and Sirt3 enhancement. A) Tissue NAD^+^ level at 4 hr post-NRH injection. N=5. B) Pan acetyl-lysine levels in liver lysates. C) Deacetylation activity of liver lysates with acetyl-p53 as substrate. N=3. D) Protein expression of SIRT3 and its downstream target, SOD2 and IDH2, and their quantification. D) Protein expression of AK, PARP1, CD38 in each group and their quantification. Data are shown as mean±SD, * indicates p<0.05, ** indicates p<0.01, *** indicates p<0.001, **** indicates p<0.0001.

ABT702 has been previously reported to degrade ADK protein after repeated administration^39,40^, however, after 7-week administration, we found no difference in hepatic ADK protein in the ABT702 or NRH treated groups (**Fig 8E**). We also assessed changes in other NAD^+^ consumer expressions, particularly PARP1 and CD38, which revealed a significant reduction in PARP1 and CD38 levels in the liver of NRH-treated mice (**Fig 8E**), indicating that NAD^+^ utilization by other consumers was suppressed, thus not competing with sirtuins for the NAD^+^ pool. ABT702 treatment alone and in combination with NRH also moderately reduced these enzyme expressions. Overall, these data suggest that although ADK inhibition did not fully block the NAD^+^ increase and sirtuin activation, it suppressed the elevation in SIRT3 protein and its downstream targets. The intact ADK function and the phosphorylation of NRH are required to transmit the full benefits of NRH in improving glucose and lipid metabolism in the DIO mice.

## Discussion

NRH is one of the most potent NAD^+^ enhancing compounds known to date. It has demonstrated higher efficacy in enhancing NAD^+^ levels both *in vitro* and *in vivo* compared to NR and NMN^21^, which are widely recognized NAD^+^ precursors showing promise in addressing obesity-related metabolic disorders^2^. In our current study, we discovered that NRH exhibited ADK-dependent activities in combating hyperglycemia and hyperlipidemia in obese, male C57BL/6J mice, further demonstrating the potential of NAD^+^ precursor in correcting metabolic diseases. In a separate ongoing study, we have observed similar treatment effects of NRH in improving glucolipid metabolism within female mice that were fed HFD.

NRH treatment was administered via IP injection three times per week, a regimen chosen because NRH sustained elevated blood NAD^+^ levels for up to 72 hours post-injection. Like other tissues, blood NAD^+^ levels are maintained through a balanced synthesis and degradation process in blood cells. NRH’s long-lasting effect on blood NAD^+^ levels reflect accelerated NAD^+^ synthesis and degradation processes throughout the body, leading to the gradual released of NAD^+^ catabolites/precursors such as nicotinamide or nicotinic acid (deamidation by intestinal microbiome^41^) into the circulation. Splanchnic tissues, including the liver, pancreas, kidney, and WAT primarily use NRH for NAD^+^ synthesis, showing a relatively transient NAD^+^ increases with peaks observed 4 hours post-treatment. At the zenith of NAD^+^ enhancement, the deacetylase activities, mainly from SIRT1-3^42^, were significantly activated. Additionally, we previously found evidence of enhanced PARP activity following NRH injection^22^. We deduce that concurrent with NRH-induced NAD^+^ surge, the activities of NAD^+^ consumers are elevated, increasing the turnover rate of NAD^+^ in the splanchnic tissues and exhausting newly synthesized NAD^+^ within 24 hour. These degradation products serve as continuous substrates for NAD^+^ synthesis in blood cells and somatic tissues such as skeletal muscle and interscapular BAT, maintaining their elevated NAD^+^ levels over 24-hour. However, administration of the ADK inhibitor, ABT702, largely hindered the utilization of NRH by splanchnic tissues, resulting in reduced NAD^+^ synthesis and diminished activation of sirtuins and PARPs. This inhibition likely also affects the subsequent release of NAD^+^ catabolites, thus reducing the supply of NAD^+^ precursors in the circulation. The unsustained blood NAD^+^ elevations and the absence of elevation at 24 hours in somatic tissues supported this theory. Therefore, the different metabolic outcomes observed with or without ADK inhibitor may be attributed to the differential activation of NAD^+^ consumer activities in visceral tissues and their subsequent release of nicotinamide or nicotinic acid into the blood, which were recycled for NAD^+^ synthesis for an extended time following the initial administration.

One of the most significant observations was that NRH exhibited specific benefits in the pancreas, inducing significant enhancements in insulin secretion and pancreatic insulin content. These findings highlighted the critical role of NAD^+^ metabolism in preserving beta cells functional mass. Previous studies have shown that depleting nicotinamide phosphoribosyltransferase (NAMPT), a key enzyme in the nicotinamide salvage pathway, reduced NAD^+^ levels and impaired GSIS response in beta cells, whereas boosting NAD^+^ levels with NMN reversed this effect^5^. Yoshino et al. reported that NMN injections improved glucose clearance rate and insulin secretion in HFD-fed or aged mice^5^. Additionally, a recent clinical trial in healthy humans showed that NMN supplementation increased postprandial plasma insulin levels^43^, further indicating a pancreatic beta cell-specific benefit of NAD^+^ enhancement. While our data agreed with the NRH’s impact in improving insulin secretion, we also showed that NRH increased total beta cell mass, which is a novel finding for NAD^+^ boosters. Although the effect of NRH on beta cell proliferation remains to be explored, our previous work demonstrated that NRH protected beta cells from hydrogen peroxide-induced cytotoxicity, suggesting a potential mechanism for preserving beta cell population by mitigating oxidative stress-induced cell death. These findings demonstrate a therapeutic potential of NRH to prevent the onset and progression of T2D in obese subjects.

The pancreatic benefits of NRH may also be related with ADK, which plays a key role in maintaining pancreatic health^38^. Knocking down or inhibiting ADK using ABT702 leads to increases in beta-cell proliferation by elevating intracellular adenosine levels and reducing DNA methylation capacities^38^. Our data showed that co-treatment of ABT702 with NRH increased beta cell mass more than either treatment alone, demonstrating a synergistic effect in promoting beta cell proliferation. It is possible that when present at large quantity, NRH competes with adenosine for the ADK binding, thereby decreasing its overall enzymatic activity and achieving effect similar to ADK inhibitor. However, this theory requires more evidence. ADK has much smaller *K*_M_ for adenosine (0.2-0.4 μM)^34^ than NRH (380 μM)^22^, and the pancreatic NAD^+^ enhancement induced by NRH was mostly transient, suggesting that any inhibitory effect may also be short-lasting. Further research is needed to understand the interaction between NRH and ADK, as well as how NRH influences DNA methylation patterns and proliferative machinery within beta cells.

One unexpected finding here was that NRH increased liver weight in DIO mice. Previous research on NAD^+^ precursors, particularly NR, has shown mixed effects on non-alcoholic fatty liver disease (NAFLD). Gariani et al. demonstrated that NR administered at 400 mg/kg/day prevented and reversed NAFLD by enhancing hepatic β-oxidation and increasing mitochondrial complex content and activity^13^. In contrast, Shi et al. reported that NR administered at a higher dose (∼1000 mg/kg/day) exacerbated metabolic dysfunction, including increasing liver weight and TG content, and WAT dysfunction^44^. They attributed these contrasting results to the use of a different mouse strain with functional nicotinamide nucleotide transhydrogenase (Nnt), which catalyzes the reduction of NADP^+^ to NADPH, compared to C57BL/6J mice that have mutated Nnt and reduced activity^45^. In our study, using C57BL/6J mice that were obese and likely had fatty livers before treatment, the mechanisms leading to increased steatosis and liver weights with NRH are likely different from those observed with high-dose NR. Specifically, NRH significantly increased liver CE accumulation and reduced circulating cholesterol levels. When an ADK inhibitor was co-treated with NRH, liver NAD^+^ remained elevated at a level comparable to NR-induced NAD^+^ increases, yet hepatic CE accumulation and serum cholesterol-lowering effects were blocked. These results suggest NRH acts through ADK to transfer excessive cholesterol from serum to the liver to be stored as CE, a relatively less toxic form^46^. However, whether NRH has anti-atherosclerotic effect remains to be validated. NRH-treated mice exhibited a significant reduction in HDL-C, the predominant lipoprotein form in mouse circulation, possibly due to increased HDL-C uptake by the liver. In humans, LDL-C is the predominant species contributing to plaque accumulation, while HDL-C is considered beneficial for reversing the process^47^. How NRH treatment affects cholesterol metabolism and liver health in other mouse models and human systems need to be evaluated in further studies.

In conclusion, this study demonstrated the therapeutic effects of NRH in treating obesity-related metabolic diseases. To our knowledge, this is the first report to reveal that NRH treatment can reach multiple tissues and produce multi-faceted benefits, including the amelioration of hyperglycemia and hyperlipidemia, as well as the enhancement of adipose tissue functionalities. Our findings highlight that the treatment effects of NRH are comparable to, and potentially more potent than, other NAD^+^ precursors such as NMN and NR. Although more safety analysis and mechanistic evaluations are needed in the future, the ability of NRH to simultaneously target and improve multiple metabolic parameters underscores its promise for future clinical applications in human metabolic disease management.

## Materials and methods

### Chemicals

All chemicals were obtained from Sigma-Aldrich if not otherwise specified. NRH was synthesized in our lab using published methods^21^. NRH used for animal administration was frequently tested with HPLC and LCMS. The purity of NRH was assessed by its peak at 340 nm, and the potential degradation products, NR and nicotinamide, were examined by their peaks at 260 nm. The NRH used in mice had over 85% purity overall.

### Animals

3-month-old, male C57BL/6J mice were purchased from the Jackson Laboratory. The animals were maintained in an animal room with controlled light (12:12 hr light-dark cycle) and temperature (21-23 °C). The animals had ad libitum access to food and water. In the first study, age-matched mice were provided with either chow or HFD for 3 months, then randomly assigned to vehicle control or NRH treatment groups. The mice received injections with sterile PBS or 250 mg/kg NRH dissolved in PBS via IP injection every Monday, Wednesday, and Friday at the same time for 7 weeks (n=5 for chow-fed groups, n=20 for HFD-fed groups). In the second study, mice were fed HFD for 3 months, then randomly assigned to vehicle control, ABT702, NRH, and NRH+ABT702 groups. These mice received injections with sterile vehicle, 3 mg/kg ABT702, 250 mg/kg NRH, or ABT702 followed by 250 mg/kg NRH after 1 hour through IP injection, every Monday, Wednesday, and Friday at the same time for 7 weeks (n=5 per group). The food consumption and body weights were recorded weekly. The mice were euthanized after 16 hours of fasting at the end of the experiment. Blood was collected through cardiac puncture, then separated for serum and uncoagulated whole blood. The liver, kidney, pancreas, heart, white adipose tissues, skeletal muscle, and brown adipose tissue were collected and weighed. Some of the tissue samples were fixed with 4% paraformaldehyde and later embedded into paraffin blocks. The rest of the tissue samples were aliquoted and snap frozen. All tissues were stored at -80 °C until analysis. All procedures were approved by the Institutional Animal Care and Use Committee of Weill Cornell Medicine.

### Blood and tissue NAD level measurement

For whole blood, 50 µL of blood was weighed, then dissolved in 50 µL 7% perchloric acid for NAD extraction. For tissues, 30-50 mg of tissues were pulverized in liquid nitrogen and homogenized in 7% perchloric acid by sonication, and then the solution was neutralized and subjected to the published cycling assay for NAD measurement according to a previously published protocol^48^. The NAD levels were normalized to the original tissue weight.

### Glucose tolerance test

To assess glucose homeostasis, all mice were fasted overnight for 16 hours with free access to water. The dextrose solution (10% in sterilized PBS) was IP injected at 2 mg/g body weight into each mouse. Tail blood from each mouse was tested using the AlphaTRAK® Glucose Meter (Zoetis) before and at 15, 30, 60, 90, and 120 minutes after injection.

### Insulin Tolerance Test

To assess insulin sensitivity, mice were fasted for 4 hours. Insulin solution (0.75 IU/kg body weight for chow-fed mice, 2 IU/kg body weight for HFD-fed mice) was IP injected into each mouse. Blood glucose levels were tested at 0, 15, 30, 45, and 90 minutes.

### Pyruvate Tolerance Test

To determine the glucose production rate, 2 mg/g body weight pyruvic acid dissolved in sterile PBS was injected after 16 hours of fasting. Blood glucose levels were measured every 15 minutes for over 2 hours.

### Glucose-stimulated insulin secretion

To measure insulin secretion, whole blood was collected at 0, 15, and 30-minute time points during the GTT challenge. Serum was isolated after letting the whole blood clot at room temperature for 30 minutes, then centrifuged at 1,000 RCF for 15 minutes at 4°C. Insulin levels were determined using a Mouse Insulin ELISA kit (Crystal Chem) following the manufacturer’s instructions.

### Pancreatic insulin extraction and quantification

Approximately 50 mg of frozen pancreas tissue was pulverized in liquid nitrogen and homogenized in an acid/ethanol solution (0.18 M HCl, 70% ethanol). The homogenates were kept overnight at 4°C before being centrifuged at 10,000 RCF for 20 minutes at 4°C, and the supernatant was used for the insulin assay with the previously described kit. The pancreatic insulin content was normalized to the total tissue weight in each sample.

### HOMA-IR calculation

The HOMA-IR was calculated using the equation [fasting plasma glucose (mmol/L) × fasting serum insulin (µU/mL) / 22.5] ^49^.

### Histological analysis

Formalin-fixed tissues were embedded in paraffin blocks using Tissue-Tek VIP® tissue processor (Sakura Finetek USA, Inc.). Paraffin sectioning and the hematoxylin and eosin (H&E) staining were performed through the Electron Microscopy & Histology Core at Weill Cornell Medicine. For immunohistochemistry (IHC) analysis, 5 μm thick pancreas tissue sections were deparaffinized in xylene and rehydrated. The tissue slides were incubated with fresh prewarmed antigen retrieval buffer (Vector Labs), and the buffer was heated to 100°C. The tissue slides were then equilibrated to room temperature for 30 min. After being added to 3% H_2_O_2_ and rinsed with PBS, the slides were incubated with antibodies against insulin at 4°C overnight. After three washes with PBS, the tissue slides were incubated with the secondary antibody (Thermo Fisher Scientific) for 30 min at room temperature. The slides were then color-reacted with 3,3’-Diaminobenzidine (DAB) (Vector Labs) and counterstained with hematoxylin. Images were taken with Zeiss Axioscope 5 microscope (Leica Microsystems Inc.).

For beta cell mass quantification, insulin-stained pancreas slides were imaged. The percentage of insulin-labeled area was quantified employing Adobe Photoshop (San Jose) and ImageJ as described ^50^.

### Tissue triglyceride (TG) measurement

Samples of liver and skeletal muscle (each weighing 50 mg) were homogenized in PBS and subsequently extracted by the Folch method^51^ with a chloroform-methanol mixture in a ratio of 2:1 (v/v). The resulting residue was then re-suspended in a solution containing 1% Triton X-100 prepared in 100% ethanol. TG content was quantified using assay kit (Thermo Fisher Scientific).

### Body Composition and Metabolic cage

Body composition, including lean and fat mass, was spectroscopy with an EchoMRI 3-in-1 Body Composition Analyzer (EchoMRI). Metabolic monitoring was performed using the Promethion Metabolic Screening System (Promethion High-Definition Multiplexed Respirometry System for Mice; Sable Systems International). Metabolic cages were housed within temperature-controlled environmental chambers (DB034 Laboratory Incubator, Darwin Chambers). Mice tested (n=5 per group) were individually caged in fully enclosed metabolic cages equipped with HEPA filters to ensure a sterile airflow and acclimated for 3 days before measurement. Within the 24 hours of measurement, the rates of oxygen consumption (VO_2_) and carbon dioxide production (VCO_2_) were measured by indirect calorimetry, with a sampling frequency set at 1 second. Respiratory data were collected every 5 minutes, with each cage being sampled for 30 seconds and a baseline cage sampling frequency of 30 seconds after every four cages. The respiratory exchange ratio was calculated as the ratio of VCO_2_ to VO_2_. Continuous measurements of food intake were taken through gravimetric methods within the metabolic cages. The distance traveled by the mice was assessed based on the number of beam breaks detected by a grid of infrared sensors integrated into each cage.

### Serum chemistry analysis

Serum concentrations of various biochemical markers, including Alanine Aminotransferase (ALT), Aspartate Aminotransferase (AST), TG, Total cholesterol, High-Density Lipoprotein Cholesterol (HDL-C), Low-Density Lipoprotein Cholesterol (LDL-C) are measured through serum chemistry service at Laboratory of Comparative Pathology at Weill Cornell Medicine. Other proteins, including adiponectin (RayBiotech), HMW adiponectin (MyBioSource), Adipsin (R&D Systems), and Interleukin-6 (Proteintech), were measured using ELISA kits. Thiobarbituric Acid Reactive Substances (TBARS) (Cayman Chemical Company) and Free fatty acid (Abcam) were measured using kits following the manufacturers’ protocols.

### Realtime qPCR

RNA was extracted from tissue samples using TRIZOL-chloroform and synthesized into cDNA with reverse transcriptase (Quantabio). The gene expression levels were determined by quantitative real-time PCR using the SYBR Green SuperMix Reagent (Quantabio). The PCR was performed in triplicate and run on a StepOnePlus Real-Time PCR System (Applied Biosystems). 18S was used as housekeeping control. The primer sequences are listed in Supplement Table 1.

### Western Blot

Proteins from mice tissue were extracted using RIPA buffer, with the addition of protease and phosphatase inhibitors cocktail, 5 μM Trichostatin A and 5 mM Nicotinamide. Protein concentrations were quantified using Bradford assay (Thermo Fisher Scientific). 30 μg of protein from each sample were electrophoresed on a 4-12% SDS gradient gels and transferred to polyvinylidene difluoride membranes (Nitrocellulose; Millipore Corp). 1% BSA was used as blocking buffer for all phosphate proteins, and 5% non-fat milk for other proteins. After for 1 h in Tris-buffered saline containing 0.1% Tween-20 (TBST), the membranes were incubated at 4°C overnight with primary antibodies. The membranes were washed with TBST for 30 min and then immunoblotted with HRP-conjugated secondary antibodies at room temperature for 1 h. Blots were developed by enhanced chemiluminescence (Thermo Fisher Scientific). Autoradiograms were densitometrical determined by using the Quantity One software package (Bio-Rad). a-tubulin and GAPDH was used as a loading control and were visualized using monoclonal antibodies against them. Western blots were analyzed using Image-Lab software (version 1.41). All antibody information can be found in Supplement Table 2.

### P53 deacetylation activity

Activity of p53 deacetylation was test by a sirtuin activity assay kit (Biotechnology Research). Briefly, the acetylated p53-AFC substrate undergoes deacetylation by the Sirtuins, releasing a fluorophore that can be detected at excitation/emission wavelengths of 400/505 nm using a plate reader (Spectrmax M4).

### Lipidomic

Liver tissues (20–30 mg) were homogenized in 100 μl of PBS buffer, then extracted with 20 volumes of chloroform-methanol (2:1) in a single-phase extraction process, recovering all lipids in a single phase suitable for liquid chromatography-mass spectrometry (LC-MS) analysis. Lipid analyses were performed using an HP 1200 LC system combined with a PE Sciex API 4000 Q/TRAP MS with a turbo-ionspray source (350°C) and Analyst 1.5 data system. Data are shown in the Log2FC in the heatmap, and p-values were adjusted using False discovery rate.

### Statistical analysis

Statistical analyses were done using Prism 9.1.0 software (GraphPad Software, Inc). Data were shown as mean ± standard deviation (SD). Comparison between two groups were performed with unpaired t-test. Multiple comparisons were performed with Ordinary one-way ANOVA. Differences were considered significant if p value is less than 0.05.

## Reference

1 Jaacks, L. M. et al. The obesity transition: stages of the global epidemic. Lancet Diabetes Endocrinol 7, 231–240, doi:10.1016/S2213-8587(19)30026-9 (2019).

2 Yang, Y. & Sauve, A. A. NAD(+) metabolism: Bioenergetics, signaling and manipulation for therapy. Biochim Biophys Acta 1864, 1787–1800, doi:10.1016/j.bbapap.2016.06.014 (2016).

3 Akasaka, H. et al. Effects of nicotinamide mononucleotide on older patients with diabetes and impaired physical performance: A prospective, placebo-controlled, double-blind study. Geriatr Gerontol Int 23, 38–43, doi:10.1111/ggi.14513 (2023).

4 Zhong, O., Wang, J., Tan, Y., Lei, X. & Tang, Z. Effects of NAD+ precursor supplementation on glucose and lipid metabolism in humans: a meta-analysis. Nutr Metab (Lond) 19, 20, doi:10.1186/s12986-022-00653-9 (2022).

5 Yoshino, J., Mills, K. F., Yoon, M. J. & Imai, S. Nicotinamide mononucleotide, a key NAD(+) intermediate, treats the pathophysiology of diet-and age-induced diabetes in mice. Cell Metab 14, 528–536, doi:10.1016/j.cmet.2011.08.014 (2011).

6 Liu, S., Kim, T. H., Franklin, D. A. & Zhang, Y. Protection against High-Fat-Diet-Induced Obesity in MDM2(C305F) Mice Due to Reduced p53 Activity and Enhanced Energy Expenditure. Cell Rep 18, 1005–1018, doi:10.1016/j.celrep.2016.12.086 (2017).

7 Gomes, A. P. et al. Declining NAD(+) induces a pseudohypoxic state disrupting nuclear-mitochondrial communication during aging. Cell 155, 1624–1638, doi:10.1016/j.cell.2013.11.037 (2013).

8 Imai, S. & Guarente, L. NAD+ and sirtuins in aging and disease. Trends Cell Biol 24, 464–471, doi:10.1016/j.tcb.2014.04.002 (2014).

9 Tarrago, M. G. et al. A Potent and Specific CD38 Inhibitor Ameliorates Age-Related Metabolic Dysfunction by Reversing Tissue NAD(+) Decline. Cell Metab 27, 1081–1095 e1010, doi:10.1016/j.cmet.2018.03.016 (2018).

10 Bai, P. et al. PARP-1 inhibition increases mitochondrial metabolism through SIRT1 activation. Cell Metab 13, 461–468, doi:10.1016/j.cmet.2011.03.004 (2011).

11 Trammell, S. A. et al. Nicotinamide Riboside Opposes Type 2 Diabetes and Neuropathy in Mice. Sci Rep 6, 26933, doi:10.1038/srep26933 (2016).

12 Canto, C. et al. The NAD(+) precursor nicotinamide riboside enhances oxidative metabolism and protects against high-fat diet-induced obesity. Cell Metab 15, 838–847, doi:10.1016/j.cmet.2012.04.022 (2012).

13 Gariani, K. et al. Eliciting the mitochondrial unfolded protein response by nicotinamide adenine dinucleotide repletion reverses fatty liver disease in mice. Hepatology 63, 1190–1204, doi:10.1002/hep.28245 (2016).

14 Revollo, J. R. et al. Nampt/PBEF/Visfatin regulates insulin secretion in beta cells as a systemic NAD biosynthetic enzyme. Cell Metab 6, 363–375, doi:10.1016/j.cmet.2007.09.003 (2007).

15 Mills, K. F. et al. Long-Term Administration of Nicotinamide Mononucleotide Mitigates Age-Associated Physiological Decline in Mice. Cell Metab 24, 795–806, doi:10.1016/j.cmet.2016.09.013 (2016).

16 Sauve, A. A. et al. Triple-Isotope Tracing for Pathway Discernment of NMN-Induced NAD(+) Biosynthesis in Whole Mice. Int J Mol Sci 24, doi:10.3390/ijms241311114 (2023).

17 Liu, L. et al. Quantitative Analysis of NAD Synthesis-Breakdown Fluxes. Cell Metab 27, 1067–1080 e1065, doi:10.1016/j.cmet.2018.03.018 (2018).

18 Kim, L. J. et al. Host-microbiome interactions in nicotinamide mononucleotide (NMN) deamidation. FEBS Lett 597, 2196–2220, doi:10.1002/1873-3468.14698 (2023).

19 Damgaard, M. V. & Treebak, J. T. What is really known about the effects of nicotinamide riboside supplementation in humans. Sci Adv 9, eadi4862, doi:10.1126/sciadv.adi4862 (2023).

20 Song, Q. et al. The Safety and Antiaging Effects of Nicotinamide Mononucleotide in Human Clinical Trials: an Update. Adv Nutr 14, 1416–1435, doi:10.1016/j.advnut.2023.08.008 (2023).

21 Yang, Y., Mohammed, F. S., Zhang, N. & Sauve, A. A. Dihydronicotinamide riboside is a potent NAD(+) concentration enhancer in vitro and in vivo. J Biol Chem 294, 9295–9307, doi:10.1074/jbc.RA118.005772 (2019).

22 Yang, Y., Zhang, N., Zhang, G. & Sauve, A. A. NRH salvage and conversion to NAD(+) requires NRH kinase activity by adenosine kinase. Nat Metab 2, 364–379, doi:10.1038/s42255-020-0194-9 (2020).

23 Williams, L. M. et al. The development of diet-induced obesity and glucose intolerance in C57BL/6 mice on a high-fat diet consists of distinct phases. PLoS One 9, e106159, doi:10.1371/journal.pone.0106159 (2014).

24 Klop, B., Elte, J. W. & Cabezas, M. C. Dyslipidemia in obesity: mechanisms and potential targets. Nutrients 5, 1218–1240, doi:10.3390/nu5041218 (2013).

25 Romani, M., Hofer, D. C., Katsyuba, E. & Auwerx, J. Niacin: an old lipid drug in a new NAD(+) dress. J Lipid Res 60, 741–746, doi:10.1194/jlr.S092007 (2019).

26 Lo, J. C. et al. Adipsin is an adipokine that improves beta cell function in diabetes. Cell 158, 41–53, doi:10.1016/j.cell.2014.06.005 (2014).

27 Zhu, N. et al. High-molecular-weight adiponectin and the risk of type 2 diabetes in the ARIC study. J Clin Endocrinol Metab 95, 5097–5104, doi:10.1210/jc.2010-0716 (2010).

28 Gan, L. T. et al. Hepatocyte free cholesterol lipotoxicity results from JNK1-mediated mitochondrial injury and is HMGB1 and TLR4-dependent. J Hepatol 61, 1376–1384, doi:10.1016/j.jhep.2014.07.024 (2014).

29 Covey, S. D., Krieger, M., Wang, W., Penman, M. & Trigatti, B. L. Scavenger receptor class B type I-mediated protection against atherosclerosis in LDL receptor-negative mice involves its expression in bone marrow-derived cells. Arterioscler Thromb Vasc Biol 23, 1589–1594, doi:10.1161/01.ATV.0000083343.19940.A0 (2003).

30 Hussain, M. M., Rava, P., Walsh, M., Rana, M. & Iqbal, J. Multiple functions of microsomal triglyceride transfer protein. Nutr Metab (Lond) 9, 14, doi:10.1186/1743-7075-9-14 (2012).

31 Stein, L. R. & Imai, S. The dynamic regulation of NAD metabolism in mitochondria. Trends Endocrinol Metab 23, 420–428, doi:10.1016/j.tem.2012.06.005 (2012).

32 Brunet, A. et al. Stress-dependent regulation of FOXO transcription factors by the SIRT1 deacetylase. Science 303, 2011–2015, doi:10.1126/science.1094637 (2004).

33 Ferber, E. C. et al. FOXO3a regulates reactive oxygen metabolism by inhibiting mitochondrial gene expression. Cell Death Differ 19, 968–979, doi:10.1038/cdd.2011.179 (2012).

34 Boison, D. Adenosine kinase: exploitation for therapeutic gain. Pharmacol Rev 65, 906–943, doi:10.1124/pr.112.006361 (2013).

35 Boison, D. et al. Neonatal hepatic steatosis by disruption of the adenosine kinase gene. Proc Natl Acad Sci U S A 99, 6985–6990, doi:10.1073/pnas.092642899 (2002).

36 Paz, J. A. et al. Adenosine kinase deficiency presenting with tortuous cervical arteries: A risk factor for recurrent stroke. JIMD Rep 62, 49–55, doi:10.1002/jmd2.12252 (2021).

37 Staufner, C. et al. Adenosine kinase deficiency: expanding the clinical spectrum and evaluating therapeutic options. J Inherit Metab Dis 39, 273–283, doi:10.1007/s10545-015-9904-y (2016).

38 Annes, J. P. et al. Adenosine kinase inhibition selectively promotes rodent and porcine islet beta-cell replication. Proc Natl Acad Sci U S A 109, 3915–3920, doi:10.1073/pnas.1201149109 (2012).

39 Fassett, J. et al. Adenosine kinase attenuates cardiomyocyte microtubule stabilization and protects against pressure overload-induced hypertrophy and LV dysfunction. J Mol Cell Cardiol 130, 49–58, doi:10.1016/j.yjmcc.2019.03.015 (2019).

40 Wolkart, G. et al. Adenosine kinase (ADK) inhibition with ABT-702 induces ADK protein degradation and a distinct form of sustained cardioprotection. Eur J Pharmacol 927, 175050, doi:10.1016/j.ejphar.2022.175050 (2022).

41 Feng, S., Guo, L., Wang, H., Yang, S. & Liu, H. Bacterial PncA improves diet-induced NAFLD in mice by enabling the transition from nicotinamide to nicotinic acid. Commun Biol 6, 235, doi:10.1038/s42003-023-04613-8 (2023).

42 Michishita, E., Park, J. Y., Burneskis, J. M., Barrett, J. C. & Horikawa, I. Evolutionarily conserved and nonconserved cellular localizations and functions of human SIRT proteins. Mol Biol Cell 16, 4623–4635, doi:10.1091/mbc.e05-01-0033 (2005).

43 Yamane, T., Imai, M., Bamba, T. & Uchiyama, S. Nicotinamide mononucleotide (NMN) intake increases plasma NMN and insulin levels in healthy subjects. Clin Nutr ESPEN 56, 83–86, doi:10.1016/j.clnesp.2023.04.031 (2023).

44 Shi, W. et al. High Dose of Dietary Nicotinamide Riboside Induces Glucose Intolerance and White Adipose Tissue Dysfunction in Mice Fed a Mildly Obesogenic Diet. Nutrients 11, doi:10.3390/nu11102439 (2019).

45 Close, A. F., Chae, H. & Jonas, J. C. The lack of functional nicotinamide nucleotide transhydrogenase only moderately contributes to the impairment of glucose tolerance and glucose-stimulated insulin secretion in C57BL/6J vs C57BL/6N mice. Diabetologia 64, 2550–2561, doi:10.1007/s00125-021-05548-7 (2021).

46 Miyazaki, M., Kim, Y. C., Gray-Keller, M. P., Attie, A. D. & Ntambi, J. M. The biosynthesis of hepatic cholesterol esters and triglycerides is impaired in mice with a disruption of the gene for stearoyl-CoA desaturase 1. J Biol Chem 275, 30132–30138, doi:10.1074/jbc.M005488200 (2000).

47 Gordon, S. M. et al. A comparison of the mouse and human lipoproteome: suitability of the mouse model for studies of human lipoproteins. J Proteome Res 14, 2686–2695, doi:10.1021/acs.jproteome.5b00213 (2015).

48 Yang, Y. & Sauve, A. A. Assays for Determination of Cellular and Mitochondrial NAD(+) and NADH Content. Methods Mol Biol 2310, 271–285, doi:10.1007/978-1-0716-1433-4_15 (2021).

49 Matthews DR, H. J., Rudenski AS, Naylor BA, Treacher DF, Turner RC. . Homeostasis model assessment: insulin resistance and beta-cell function from fasting plasma glucose and insulin concentrations in man. Diabetologia. 1985 Jul;28(7):412-9. , doi:10.1007/BF00280883. (1985).

50 Pascoe, J. et al. Free fatty acids block glucose-induced beta-cell proliferation in mice by inducing cell cycle inhibitors p16 and p18. Diabetes 61, 632–641, doi:10.2337/db11-0991 (2012).

51 Folch, J., Lees, M. & Sloane Stanley, G. H. A simple method for the isolation and purification of total lipides from animal tissues. J Biol Chem 226, 497–509 (1957).

