## Supplement Table 1 for "NRH, a potent NAD+ booster, improves glucose homeostasis and lipid metabolism in diet-induced obese mice though an active adenosine kinase pathway"

**Supplement Figures and Legends**

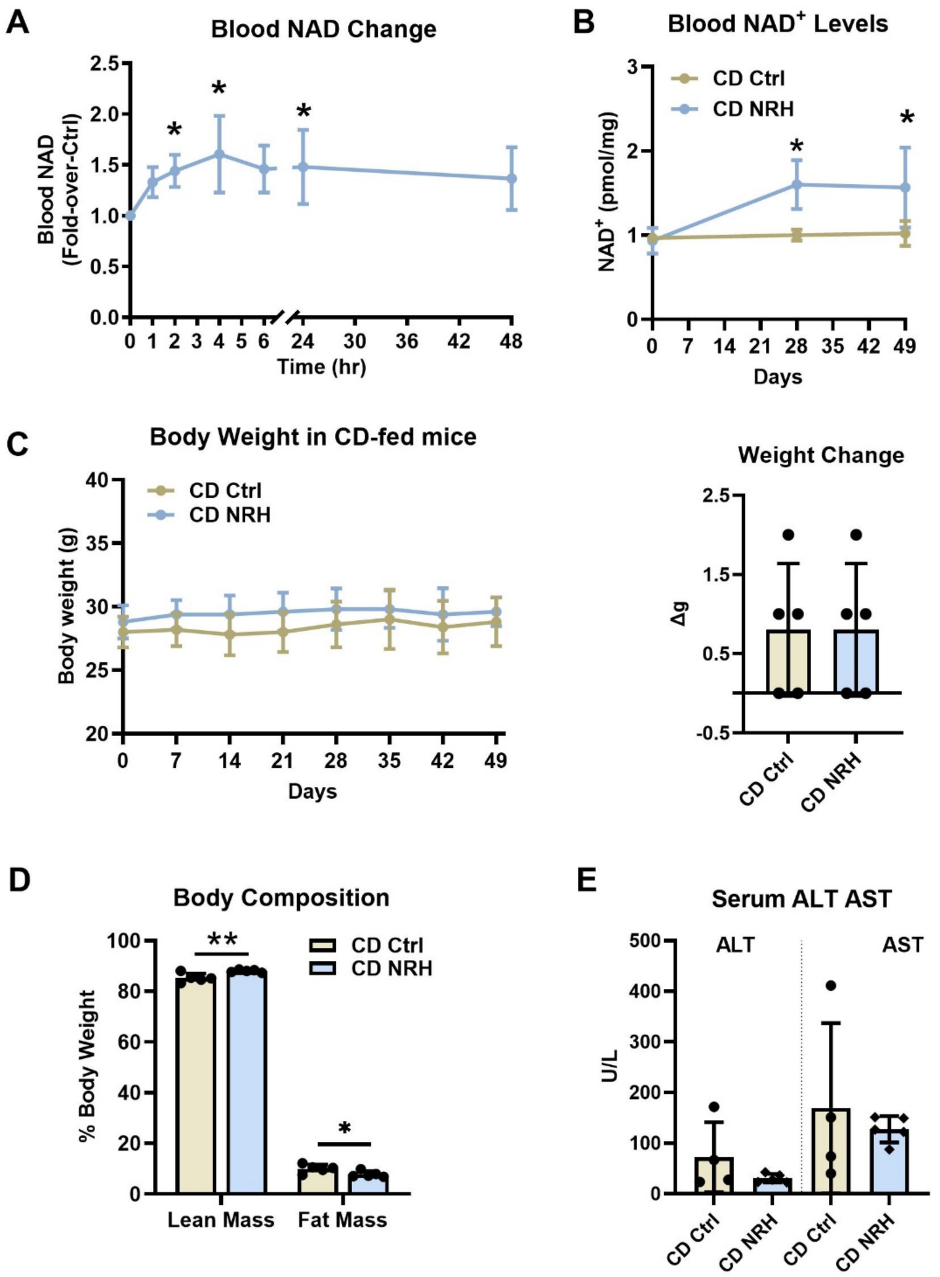


**Supplement Figure 1. NRH raised blood $NAD^+$ throughout the treatment and altering body composition in lean mice.** A) Blood $NAD^+$ levels over the first 48 hours. B) Baseline blood $NAD^+$ level throughout the 7-week treatment period. C) CD fed mice were given 250 mg/kg NRH through IP injection 3 times a week. Their body weights were recorded weekly. Total body weight change over the treatment period were calculated by subtracting the initial body weight from the final body weight. D) Body composition measurement with EchoMRI in CD-fed mice at the 5$^{th}$ week of treatment. D) Serum ALT and AST levels of CD-fed mice at sacrifice. N=5 per group. Data are shown as mean ± SD. * indicates $p<0.05$ and ** indicates $p<0.01$ compared to CD Ctrl.

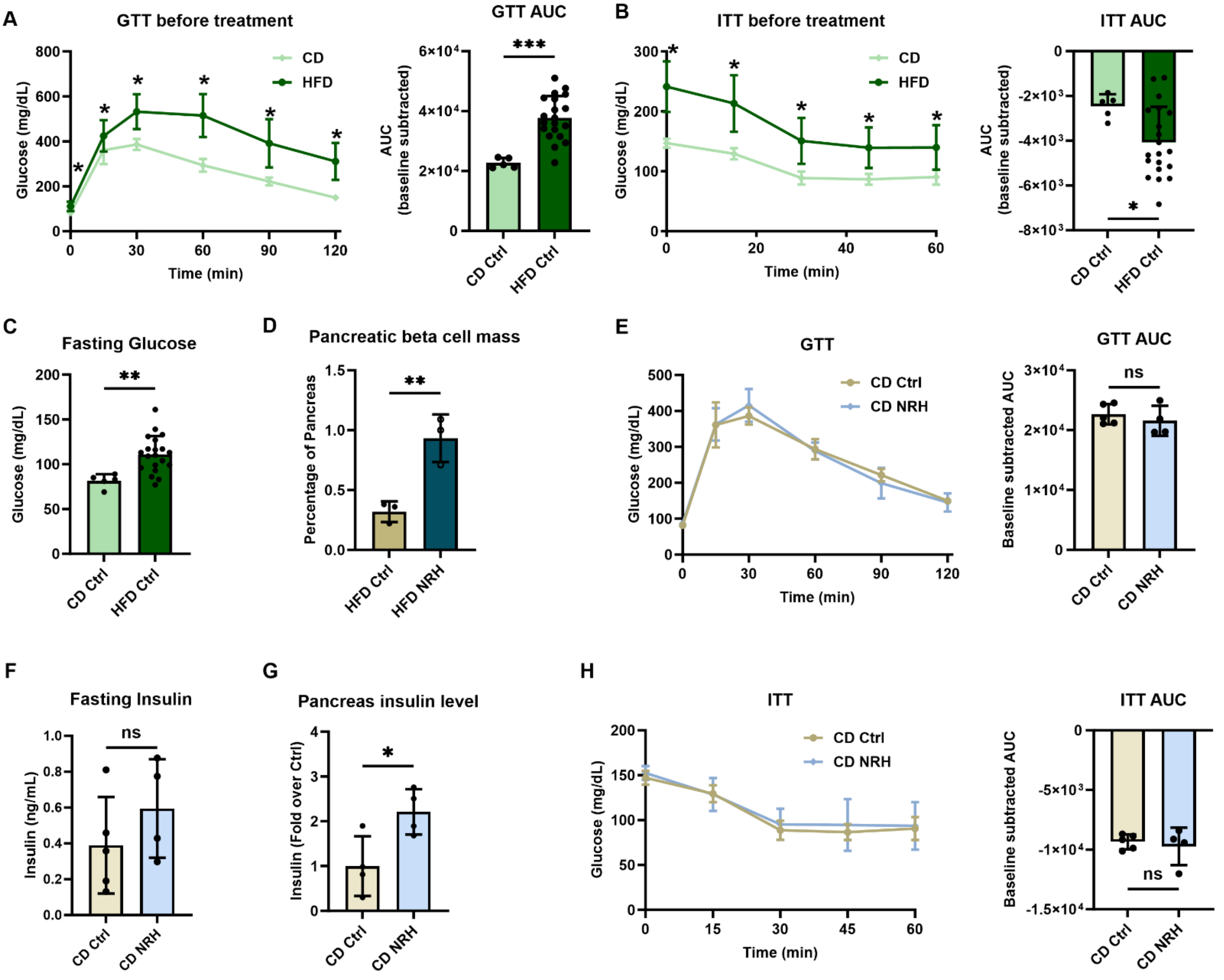


**Supplement Figure 2. NRH did not alter glucose metabolism in lean mice.** A) Comparison of GTT and their AUCs in CD-fed and HFD-fed mice before NRH treatment. B) Comparison of ITT and their AUCs in CD-fed and HFD-fed mice before NRH treatment. C) Fasting glucose in CD-fed and HFD-fed mice before NRH treatment after 16-hout fasting. D) Quantification of pancreatic beta cell mass in HFD-fed mice. E) GTT in CD-fed mice at 3rd week of treatment. F) Fasting insulin level in CD-fed mice. G) Total pancreatic insulin levels in CD-fed mice after NRH treatment. H) ITT in CD-fed mice at 6th week of treatment. N=5 in CD group and N=20 in HFD group. Data are shown as mean±SD, * indicates $p<0.05$, ** indicates $p<0.01$.

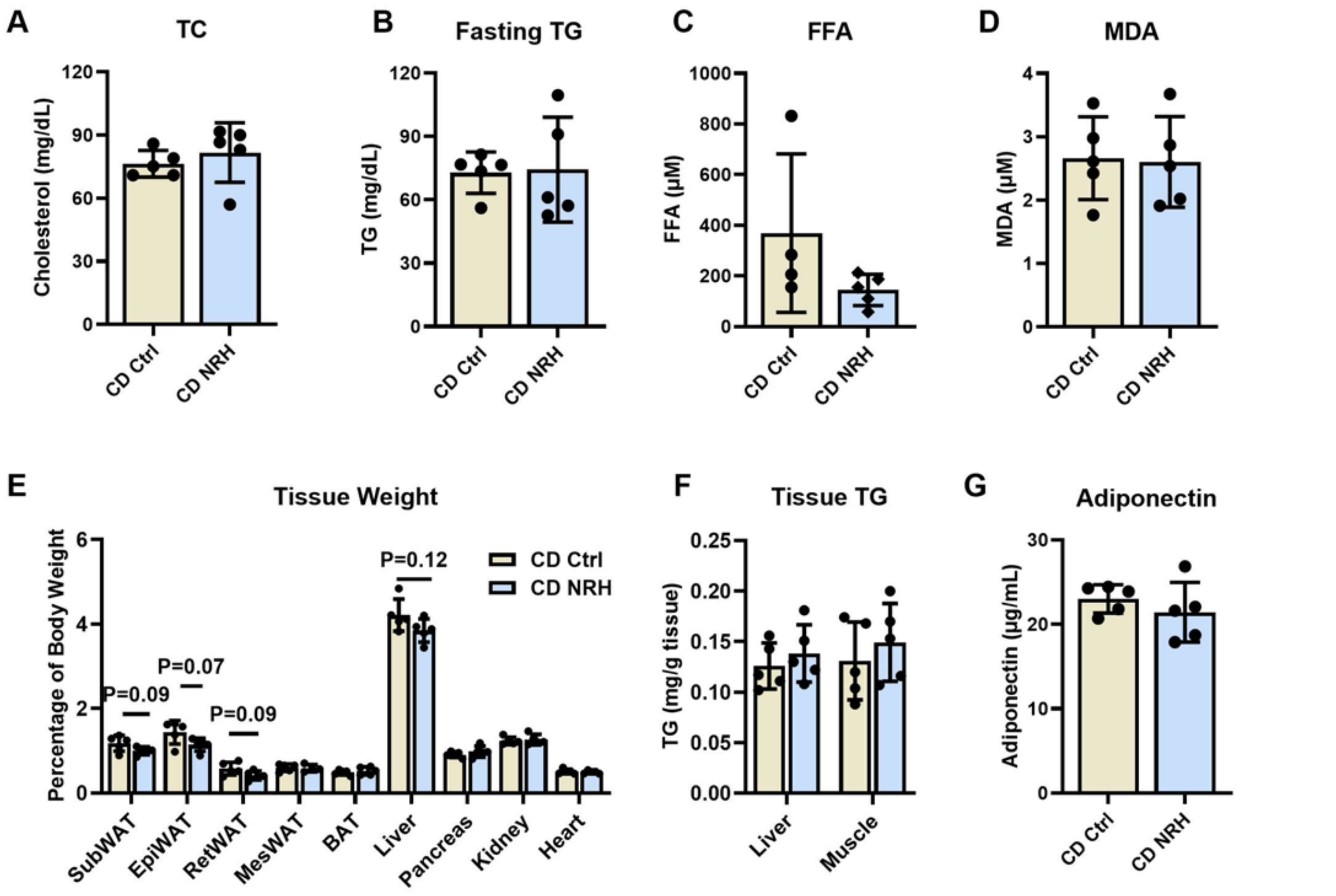


**Supplement Figure 3. NRH did not change lipid metabolism or tissue lipid deposition in lean mice.** A) Serum TC in CD-fed mice. B) Serum TG levels in 16-hour fasted lean mice. C) Serum FFA in CD-fed mice. D) Serum MDA levels in CD-fed mice. E) Tissue weights in percentage of body weight between CD-fed control and NRH treated mice. F) Tissue TG content in liver and muscle in CD-fed mice. G) Serum adiponectin levels in CD-fed mice. N=5 per group. Data are shown as mean ± SD.

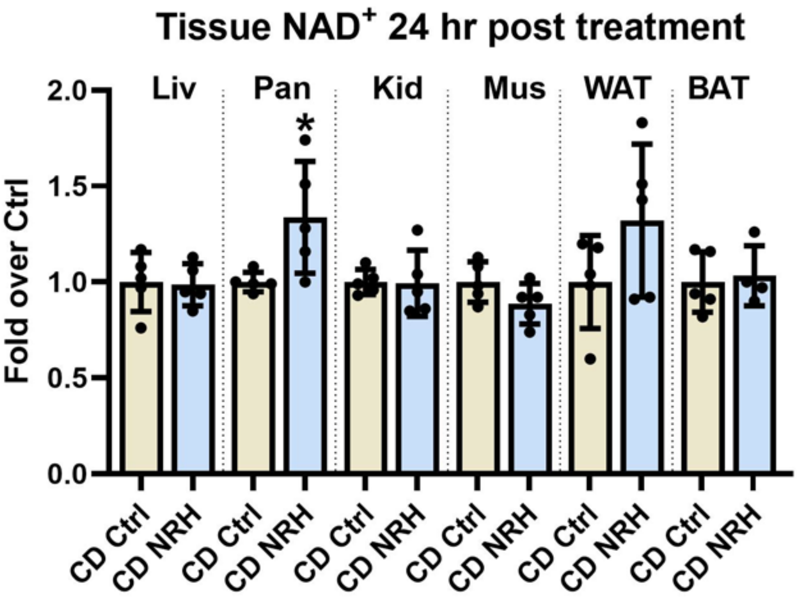


**Supplement Figure 4. NAD$^+$ changes in tissues 24 hours after NRH injection in CD-fed mice.** Data are shown as mean ± SD. N=5 per group. * indicates $p<0.05$ compared to controls.

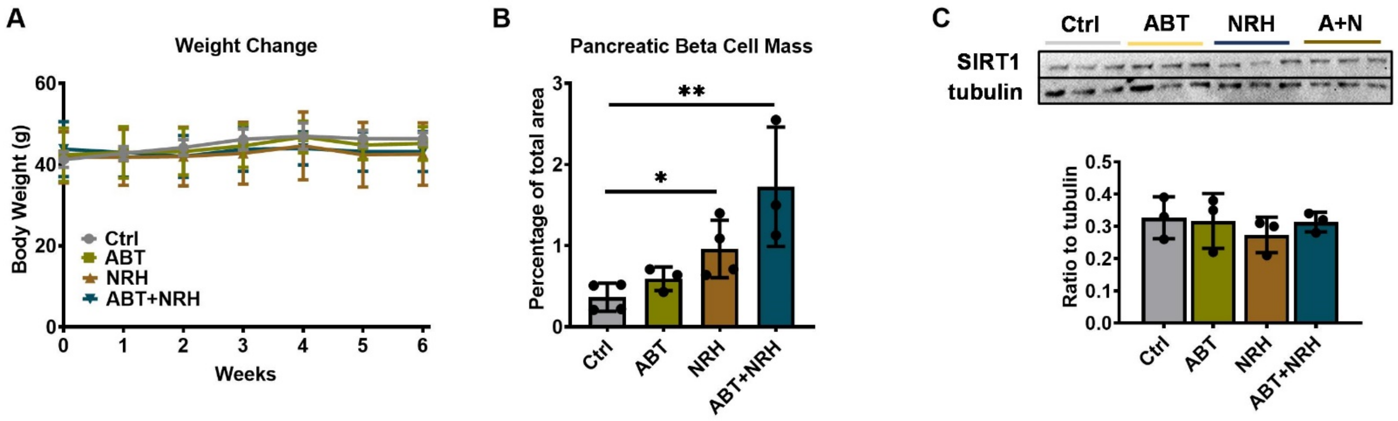


**Supplement Figure 5. Additional effects of NRH and ABT702 treatment in HFD-fed mice.** A) Average group weights in HFD-fed mice administered with different treatments over 7-week. N=5 per group. B) Ratiometric quantification of beta cell mass in the pancreas of HFD-fed mice. N=3-4 per group. C) Protein expression and quantification of SIRT1 and tubulin in the liver of HFD mice with different treatment. Data are shown as mean ± SD, * indicates $p<0.05$.

**Supplement Table1.** Realtime-PCR primer sequences

| Gene name | Forward/Reverse | Sequence |
|---|---|---|
| Srebf2 | Forward | TCTGAGGAAGGCCATTGATTAC |
| | Reverse | CAGGAAGGTGAGGACACATAAG |
| Hmgcr | Forward | CCAGAAGCTTTCGTCAGTAGAG |
| | Reverse | GCTCCCATCACCAAGGAATAA |
| Ldlr | Forward | GAGGTTCCTGTCCATCTTCTTC |
| | Reverse | GCTTCTGGGACAGCCTATTT |
| Scarb1 | Forward | GCTCAAGAATGTCCGCATAGA |
| | Reverse | AGGAATGCTCCTTTGGGTTAG |
| ApoB | Forward | GGGACTGTCTGACTTCCATATTC |
| | Reverse | CTCTCACAAGACAGGCCATATT |
| Mttp | Forward | GGCTATACAAGCTCACGTACTC |
| | Reverse | GGTGACAAAGCTGTCCTCTATC |
| Adipq | Forward | GGAAGGAGGTGGAATTGATGT |
| | Reverse | GGCGATCGAGTAGAGAAGATTG |
| Cfd | Forward | GTAAGAGGCAGGAGTCCATAAA |
| | Reverse | AACGAGGCATTCTGGGATAG |
| Lep | Forward | TGAGGGTAGAGGATGTGTTAGA |
| | Reverse | GATGCTAATGTGCCCTGAAATG |
| Pparg | Forward | CTGGCCTCCCTGATGAATAAAG |
| | Reverse | GCGGTCTCCACTGAGAATAATG |
| Plin2 | Forward | CTTGTGTCCTCCGCTTATGT |
| | Reverse | TCACTGCTCCTTTGGTCTTATC |
| Mest | Forward | CAACCTGCTGTCTCACGATTA |
| | Reverse | GCTGCCTGATTCCTGTAGTT |
| Adgre1 | Forward | GTCATCTCCCTGGTATGTCTTG |
| | Reverse | CCTCGTGTCCTTGAGTTTAGAG |
| Il6 | Forward | CTTCCATCCAGTTGCCTTCT |
| | Reverse | CCTTCTGTGACTCCAGCTTATC |
| Tnfα | Forward | GTCTCAGAATGAGGCTGGATAAG |
| | Reverse | GAGCAGAGGTTCAGTGATGTAG |
| 18s | Forward | ACGTCTGCCCTATCAACTTTC |
| | Reverse | CCGCGGTCCTATTCCATTATT |

**Supplement Table 2. Antibodies**

| | |
|---|---|
| Antibody used | Acetylated-lysine (1:1000 dilution; cat NO. 58315, Cell Signaling Technology), PCK1(1:1000 dilution; cat NO. 12940S, Cell Signaling Technology), Sirt1 (1:1000 dilution; cat NO. 9475S, Cell Signaling Technology), Sirt3 (1:1000 dilution; cat NO. 26275, Cell Signaling Technology), SOD2(1:1000 dilution; cat NO. 13141S, Cell Signaling Technology), IDH2(1:1000 dilution; cat NO. 12652S, Cell Signaling Technology), AKT (1:1000 dilution; cat NO. 9272, Cell Signaling Technology), P-AKT (1:1000 dilution; cat NO. 9271, Cell Signaling Technology), CD38(1:1000 dilution; cat NO. 51000S, Cell Signaling Technology), Parp1(1:1000 dilution; cat NO. 9532S, Cell Signaling Technology), AK (1:1000 dilution; cat NO. PA5-27399,Thermo Fisher Scientific), GAPDH (1:1000 dilution; cat NO. 2118, Cell Signaling Technology) tubulin (1:1000 dilution; cat NO. 3873, Cell Signaling Technology), insulin (1:300 dilution; cat NO. 4590, Cell Signaling Technology), anti-rabbit IgG, HRP-linked antibody (1:5000 dilution; cat NO. 7074S, Cell Signaling Technology) and anti-mouse IgG, HRP-linked antibody (1:5000 dilution; cat NO. 7076S, Cell Signaling Technology). |
| Validation | Acetylated-lysine (1:1000 dilution; cat NO. 9441, Cell Signaling Technology), Western Blot. https://www.cellsignal.com/products/primary-antibodies/acetylated-lysine-antibody/9441 |
| | PCK1(1:1000 dilution; cat NO. 12940, Cell Signaling Technology), Western Blot. https://www.cellsignal.com/products/primary-antibodies/pck1-d12f5-rabbit-mab/12940 |
| | Sirt1 (1:1000 dilution; cat NO. 9475, Cell Signaling Technology), Western Blot. https://www.cellsignal.com/products/primary-antibodies/sirt1-d1d7-rabbit-mab/9475 |
| | Sirt3 (1:1000 dilution; cat NO. 2627, Cell Signaling Technology), Western Blot. https://cellsignal.com/products/primary-antibodies/sirt3-c73e3-rabbit-mab/2627 |
| | SOD2(1:1000 dilution; cat NO. 13141, Cell Signaling Technology), Western Blot. https://www.cellsignal.com/products/primary-antibodies/sod2-d3x8f-xp-rabbit-mab/13141 |
| | IDH2(1:1000 dilution; cat NO. 12652, Cell Signaling Technology), Western Blot. https://www.cellsignal.com/products/primary-antibodies/idh2-d7h6q-rabbit-mab/12652 |
| | AKT (1:1000 dilution; cat NO. 9272, Cell Signaling Technology), Western Blot. https://www.cellsignal.com/products/primary-antibodies/akt-antibody/9272 |
| | P-AKT (1:1000 dilution; cat NO. 9271, Cell Signaling Technology), Western Blot. |

| | |
|---|---|
| | https://www.cellsignal.com/products/primary-antibodies/phospho-akt-ser473-antibody/9271 |
| | CD38(1:1000 dilution; cat NO. 51000, Cell Signaling Technology), Western Blot. https://www.cellsignal.com/products/primary-antibodies/cd38-e7z8c-xp-rabbit-mab/51000 |
| | Parp1(1:1000 dilution; cat NO. 9532, Cell Signaling Technology), Western Blot. https://www.cellsignal.com/products/primary-antibodies/parp-46d11-rabbit-mab/9532 |
| | AK (1:1000 dilution; cat NO. PA5-27399, Thermo Fisher Scientific), Western Blot. https://www.thermofisher.com/antibody/product/Adenosine-Kinase-Antibody-Polyclonal/PA5-27399 |
| | GAPDH (1:1000 dilution; cat NO. 2118, Cell Signaling Technology), Western Blot. https://www.cellsignal.com/products/primary-antibodies/gapdh-14c10-rabbit-mab/2118 |
| | tubulin (1:1000 dilution; cat NO. 3873, Cell Signaling Technology), Western Blot. https://www.cellsignal.com/products/primary-antibodies/a-tubulin-dm1a-mouse-mab/3873 |
| | insulin (1:300 dilution; cat NO. 4590, Cell Signaling Technology), Immunohistochemistry. https://www.cellsignal.com/products/primary-antibodies/insulin-antibody/4590 |
| | anti-rabbit IgG, HRP-linked antibody (1:5000 dilution; cat NO. 7074, Cell Signaling Technology), Western Blot. https://www.cellsignal.com/products/secondary-antibodies/anti-rabbit-igg-hrp-linked-antibody/7074 |
| | anti-mouse IgG, HRP-linked antibody (1:5000 dilution; cat NO. 7076, Cell Signaling Technology), Western Blot. https://www.cellsignal.com/products/secondary-antibodies/anti-mouse-igg-hrp-linked-antibody/7076 |
